# Hierarchical dynamic coding coordinates speech comprehension in the human brain

**DOI:** 10.1101/2024.04.19.590280

**Authors:** Laura Gwilliams, Alec Marantz, David Poeppel, Jean-Remi King

## Abstract

Speech comprehension involves transforming an acoustic waveform into meaning. To do so, the human brain generates a hierarchy of features that converts the sensory input into increasingly abstract language properties. However, little is known about how rapid incoming sequences of hierarchical features are continuously coordinated. Here, we propose that each language feature is supported by a dynamic neural code, which represents the sequence history of hierarchical features in parallel. To test this ‘Hierarchical Dynamic Coding’ (HDC) hypothesis, we use time-resolved decoding of brain activity to track the construction, maintenance, and update of a comprehensive hierarchy of language features spanning phonetic, word form, lexical-syntactic, syntactic and semantic representations. For this, we recorded 21 native English participants with magnetoencephalography (MEG), while they listened to two hours of short stories in English. Our analyses reveal three main findings. First, the brain represents and simultaneously maintains a sequence of hierarchical features. Second, the duration of these representations depends on their level in the language hierarchy. Third, each representation is maintained by a dynamic neural code, which evolves at a speed commensurate with its corresponding linguistic level. This HDC preserves the maintenance of information over time while limiting destructive interference between successive features. Overall, HDC reveals how the human brain maintains and updates the continuously unfolding language hierarchy during natural speech comprehension, thereby anchoring linguistic theories to their biological implementations.

## Introduction

How the human brain rapidly and robustly extracts meaning from acoustic signals during speech comprehension remains a fundamental question in neuroscience. At the level of neural representation, evidence suggests that the brain transforms the sensory input into a hierarchical set of language features, which span from speech sounds to meaning ^1^.

One body of work has studied the spatial localization of this feature hierarchy using functional Magnetic Resonance Imaging (fMRI). Phonetic ^2,3^, syllabic ^4,5^ and lexical features ^6–8^ and associated syntactic structure ^9–12^ are represented in the temporal, parietal and prefrontal cortices, with more abstract linguistic representations encoded in more distributed and higher-level activation patterns ^13–15^.

The dynamics of speech processing have been studied in a complementary body of work using electro-encephalography (EEG). Auditory responses to onsets, offsets, and fluctuations in loudness are associated with the N100 component, which peaks at approximately 100 ms^16^. Surprisal associated with phonological input–for example if a phoneme violates a task-induced phonological expectation–is associated with amplitude modulations 250-300 ms ^17–19^. This is referred to as the Phonological Mapping Negativity (PMN). Processing of the abstract “word-form” has been linked to responses across left frontal electrodes around 350 ms, through amplitude modulations associated with word frequency^20^, and cross-modal word-fragment priming^21–23^. This is referred to as the P350. Semantic manipulations, such as semantic priming and effects of context, also modulate neural activity also around 350 ms, but with an opposing polarity–the N400^24^. Finally, complexity or violations in syntactic structure is associated with a broad posterior positive deflection ∼600 ms after word presentation–the P600^25–28^. Another body of work using scalp EEG, intracranial EEG, or magneto-encephalography, has also begun to study multiple features in parallel, finding simultaneous encoding of language properties^8,29–33^.

These two literatures provide a thorough picture of response dynamics when predictions at a given level of the language hierarchy are compared to the actual input at that level, to assess the degree of (mis)match. They also demonstrate the feasibility of using continuous natural speech to study the processing of multiple features in parallel. What they do not address, however, is for how long different kinds of information are maintained, or how maintenance and integration across the hierarchy from sound to meaning are coordinated.

Due to the compositional nature of language structure–that is, phonemes combine to make syllables, which combine to make words and phrases–hierarchical processing entails integrating information over variable and nested timescales to resolve feature identity^34–36^. For example, evidence from behavioural reaction time studies and eye-tracking visual world paradigms has shown that listeners maintain lower-order information of speech for multiple seconds in the future, using them to refine lexical, syntactic, and semantic interpretations^37–39^.

The longevity of language feature encoding has the clear computational advantage of enabling the composition and resolution of higher order structures^40^. However, it is difficult to reconcile this algorithmic finding with the current dominant position that there exists a one-to-one correspondence between a given language feature and its neural representation in space and time. This account fails to explain how the brain *maintains* low-level elements long enough to integrate them into more complex units^6^, while continuously *updating* each element to keep up with the continuously unfolding speech stream to appropriately process new incoming information^34,41,42^.

These constraints theoretically apply across all levels of the hierarchy: from assembling phonemes into words, to assembling words into sentences. A new computational framework with an updated view of neural implementation is thus essential to account for how the cortex simultaneously *maintains* and *updates* each of the representations of language to build increasingly high-level representations.

Time-resolved decoding of brain activity may provide a promising tool to resolve this issue ^43–45^. By decoding the representations at each point in time, acoustic-phonetic ^3^ and visual features ^46^ have recently been shown to be embedded in a dynamic neural code. For example, in Gwilliams et al., 2022, we provided evidence for a dynamic neural code that supports acoustic-phonetic processing in speech, but it remains fully unexplored whether a dynamic code underlies more abstract feature processing, and how the parameters of that code adjust as a function of hierarchical level (Figure 1). We test whether this dynamic coding can be applied hierarchically to both maintain and update the many representations of language, while avoiding interference across successive phonemes, syllables, and words; henceforth referred to as Hierarchical Dynamic Coding (HDC). We focus on representations at the level of words and sentences, rather than at the level of discourse.

**Figure 1.**
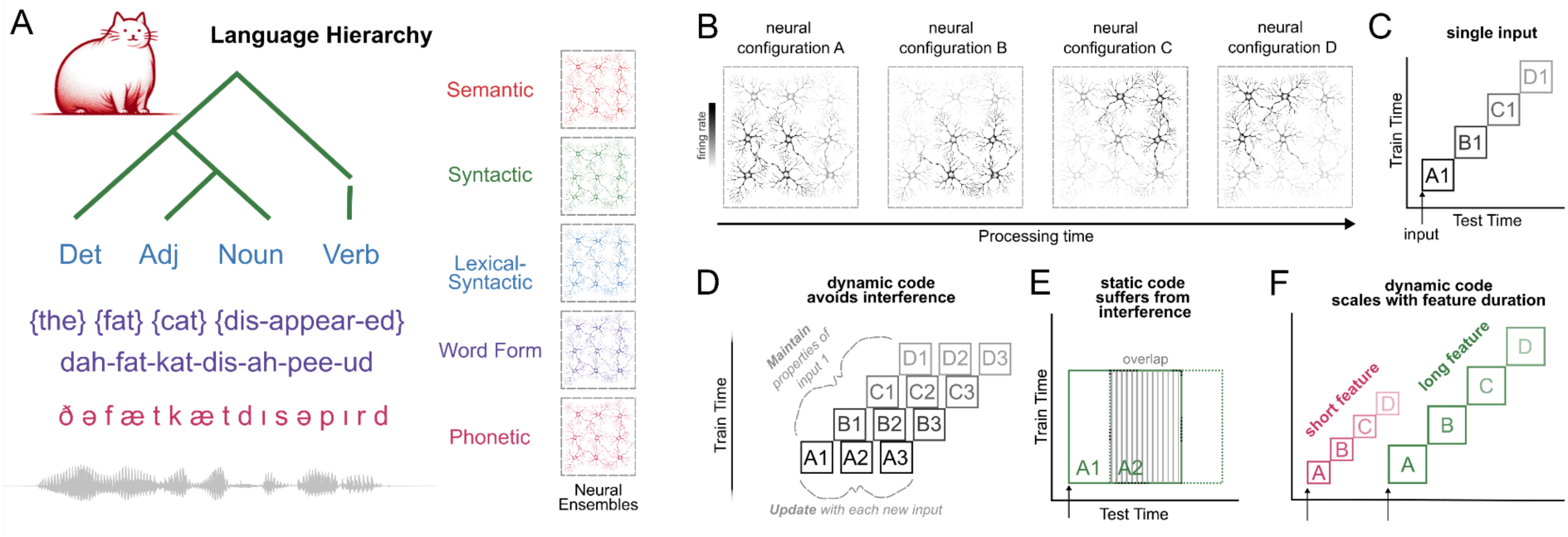
The Hierarchical Dynamic Coding Hypothesis. A: Schematic of the language hierarchy, from the acoustic input to the meaning of the utterance. Note that the image at the top of the hierarchy is intended to represent the meaning of the entire sentence, not just the single word “cat”.Each feature of the hierarchy is hypothesized to be encoded by a distinct neural ensemble. B: Schematic of the HDC hypothesis: for each feature of the hierarchy, encoding evolves across different neural ensembles as a function of time. C: Schematic decoding result for a single speech input. Each neural configuration {A, B, C, D} is engaged in sequence, leading to a lack of generalization across all train/test times. D: Schematic decoding result for sequences of inputs, which satisfies both the constraint to maintain information over time, and update as new inputs are received. This means that there is no representational overlap between neighbours in the sequence. E: A static neural code, by contrast, implies a high degree of representational overlap between neighbours. E: Schematic prediction that shorter and longer features of language will display distinct processing dynamics: shorter features will evolve between neural codes faster, and will be encoded for shorter duration; longer features will evolve slower and will be encoded for a longer duration.

To put the HDC hypothesis to the test, we recorded magneto-encephalography (MEG) from 21 participants listening to two hours of audio stories. The data used here partially overlap with the data used in Gwilliams et al., 2022 (see Methods for details). All participants were native English speakers, and the audiobooks were presented in English. We fit linear models ^43^ to decode a comprehensive set of 54 linguistic features organized into six levels of representation: phonetic, word form, lexical-syntactic, syntactic operation, semantic and syntactic state. We address three main questions: (i) can we simultaneously decode all six levels of the language hierarchy during continuous speech processing? (ii) what are the relative onsets and durations of these hierarchical levels? and (iii) does their underlying neural code evolve over time, with speed commensurate to their level in the hierarchy (Figure 1)?

## Results

### 1.1 Robust decoding of speech features

Our first question is whether the rich suite of linguistic features can be simultaneously decoded from MEG activity during continuous listening. To evaluate this, we compute the time course of each linguistic feature (Figure 2) and evaluate statistical significance with a temporal permutation cluster test of the distribution of beta coefficients across participants. Overall, our results show that we can precisely track a remarkably diverse set of linguistic features from MEG activity (Figure 3). The results of the full statistical analyses on all features are provided in Supplementary Tables 1-12. For the analysis on raw Spearman R correlation rather than B2B regression, see Supplement.

**Figure 2:**
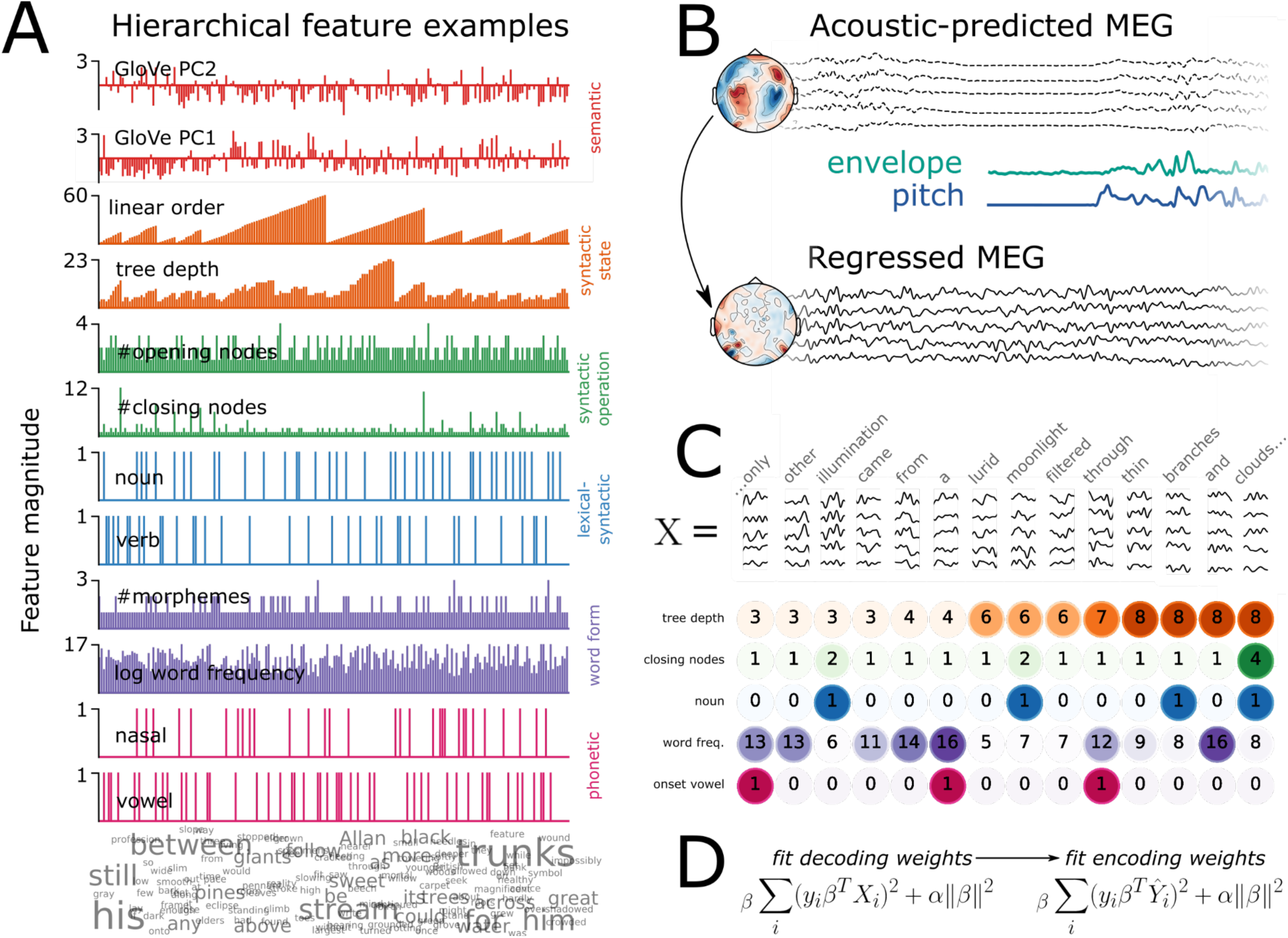
Methods. A: Feature values are plotted for a selection of 500 consecutive words, selecting two example features for each level (from a total of 54 features). The level is displayed on the right hand side. Colour corresponds to the language level it operationalises. The word cloud at the bottom displays each of the 500 words, adjusting the fontsize to be proportional to the number of instances in the story snippet. Note that the GloVe vectors are computed using a symmetric context window of size 10, and therefore they capture meaning beyond the single lexical item. B: First we fit a receptive field model based on the envelope and pitch of the acoustic speech input, and regress this out of the continuous MEG signal. The topography above shows sensor weights sensitive to acoustic features; the topography below confirms that sensor weights sensitive to acoustic featuresare at zero after this procedure. The timecourses above correspond to the TRF predictions from the acoustic model; the teal and purple timecourses correspond to the timecourse of the pitch values and envelope values; the bottom timecourse corresponds to the residual MEG data after the acoustic predictions have been subtracted out. C: Data structure. The epoch data matrix *X* has shape words (8,000) × sensors (208) × time (201). Here a schematic of the epochs is shown for a subset of the story. Below, a sample of 5 example features are displayed for each word. The superimposed number and color intensity corresponds to the feature value at a particular word. D: Main equations for the back-to-back regression method. α = the fitted regularization parameter. β = the fitted model coefficients of interest. First we fit ridge regression to decode each feature *y* from the MEG signal *X*, then we evaluate the prediction of the features (*Y*^) against the true value of *y* with an encoding model.

**Figure 3:**
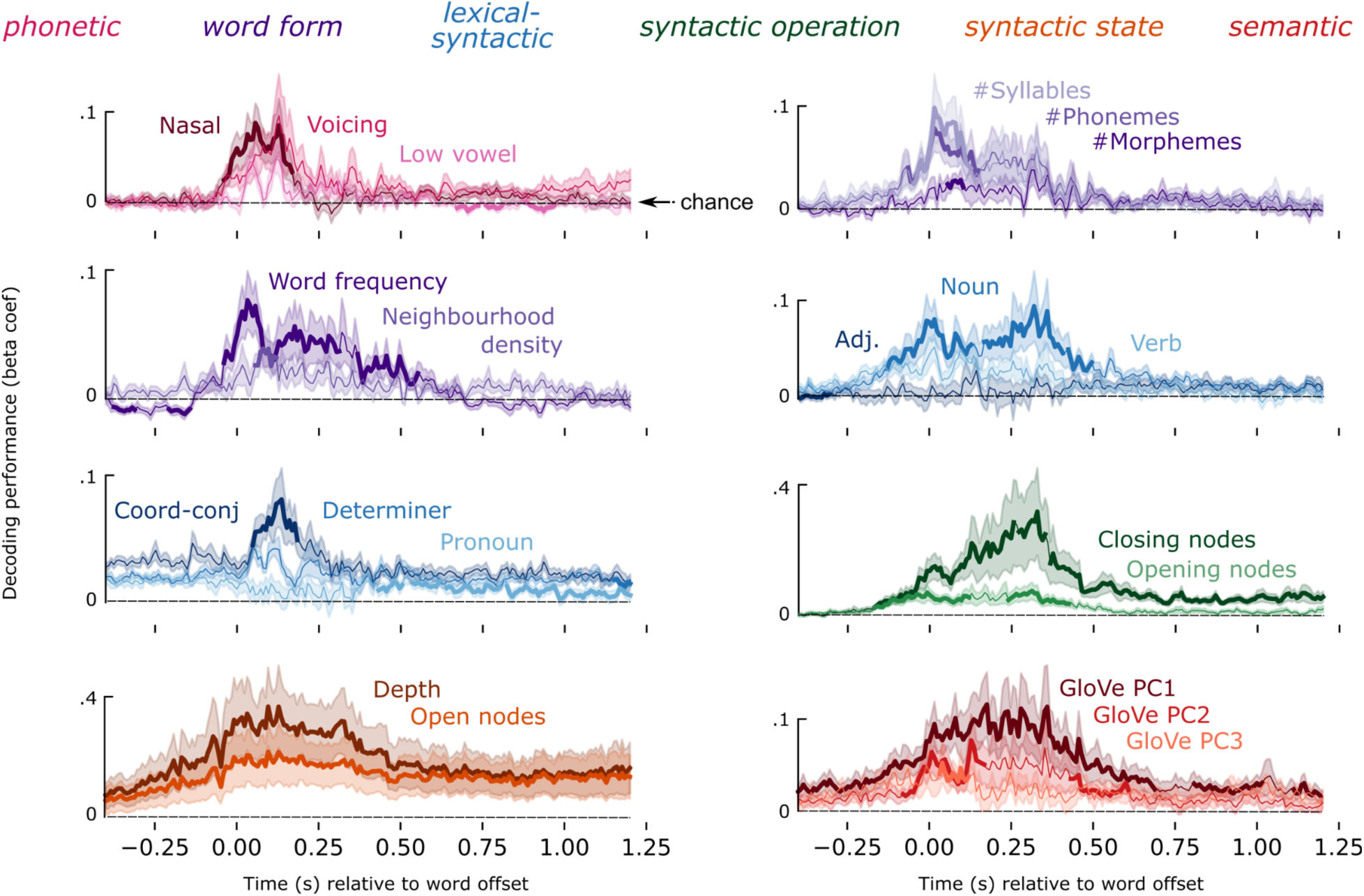
Feature decoding timecourses. Timecourses of decoding performance for a subset of language properties, locked to word offset. Line color corresponds to family assignment, which are listed above. Solid trace corresponds to mean performance across subjects; shading is the standard error of the mean across subjects. A bold mean trace corresponds to the result of a temporal permutation cluster test, indicating when the feature is decodable significantly better than chance. Dashed black line corresponds to chance-level performance.

In all results that follow, we time-lock our analyses to word offset rather than word onset, because it provides significantly higher decoding performance in all of our analyses. If the reader is interested in the word onset-locked results, or the comparison between the two, please refer to Supplementary Results.

To summarize the levels of the hierarchy that are robustly encoded in neural activity, we grouped decoding performance of the original 54 linguistic features into 6 feature families, and took the average of the features in a family, to plot 6 decoding time-courses (Figure 4A). Features across all six feature families could be detected from MEG responses, with notable differences in latency and duration: on average, phonetic features were detectable from −40:230 ms (*t*^ (average *t*-value in the cluster) = 2.57, *p* = .013) relative to word offset; word form features from – 130:550 ms (*t*^ = 2.6, *p* = .002); lexical-syntactic from −170:200 ms (*t*^ = 2.18, *p* = .029); syntactic operation from −190:1200 ms (*t*^ = 2.7, *p* < .001); syntactic state (*t*^ = 3.54, *p* < .001) and semantic vectors (*t*^ = 3.46, *p* < .001) throughout the entire search window (Figure 4A). Note that the GloVe vectors are trained using a symmetric context window of size 10, and therefore they capture meaning beyond the single lexical item. These results are consistent across the two recording sessions (Supplementary Figure 6), thus demonstrating internal replicability (see Supplement for detailed results). Overall, this analysis confirms that, during continuous speech listening, the brain builds a rich set of hierarchical linguistically motivated features.

**Figure 4:**
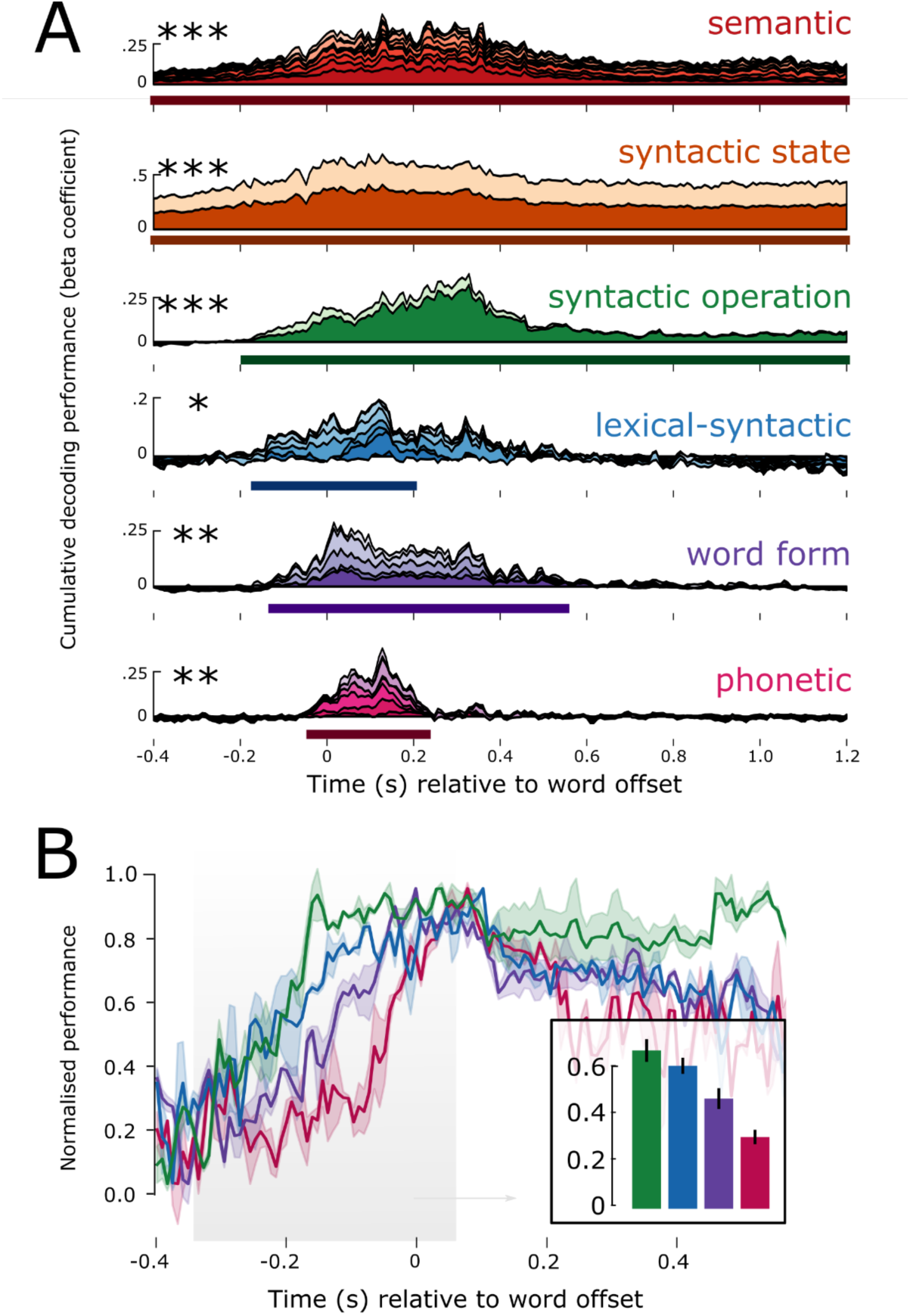
Decoding hierarchical features. A: Result of decoding each language level over time. The beta coefficients of each feature are stacked on top of each other, such that the top of the timecourse plot corresponds to the cumulative sum of all features in that linguistic level. The x-axis corresponds to time in seconds relative to word offset. The y-axis corresponds to the cumulative beta-coefficient across features. Solid line below the time-course represents the extent of the significant temporal cluster; asterisks represent its significance. B: Decoding performance zooming in for the lowest four feature families, and showing the standard error across subjects. Higher level features come online earlier than lower level ones. This is shown in the barplot, averaging performance before word offset shows a linear decrease in amplitude. * p < .05; ** p < .01; *** p < .001

### 1.2. The timing of linguistic representations depends on their level in the language hierarchy

How do the latency and duration of each feature relate to their respective level in the linguistic hierarchy? To address this issue, we analyzed the average time-course of each of the 6 feature families (Figure 4).

First, we assessed the relationship between hierarchy and decoding onset time. For this, we normalized the decoding performance for each feature family, by dividing the group average by its maximum, for each family separately. We analyzed the rise-time before word offset (for the analysis on word onset see Supplementary Figures). As shown in Figure 4B, higher-level features were detectable earlier than lower-level features, resulting in a significant negative correlation between hierarchical level and the peak of the normalized performance (*r* = −0.82, *p* < .001).

Second, we tested the relationship between hierarchy and decoding duration. We found that higher-level features were decodable significantly longer than lower-level features, resulting in a significant positive correlation between level and duration (*r* = +0.75, *p* < .001). This effect was particularly striking for the syntactic and semantic features, which were decodable for over 1s after word offset, continuing well into the processing of the subsequent words (see Supplementary Figure 1 for the distribution of latencies of upcoming words).

Third, we tested the extent to which different features of the hierarchy are represented in parallel. We found evidence of a nested temporal structure, whereby the decodable window of a given level (L) was generally contained within the decodable window of the feature at L+1. For example, the start and end of significant phonetic decoding falls within the start and end of word form decoding, and that in turn within the start and end of lexical-syntactic decoding, etc. A one-way F-test revealed that the entire hierarchy as defined by the 6 feature families was decodable in parallel from −40:230 ms (*F*-value in the cluster) = 4.1, *p* < .001) relative to word offset, i.e., throughout the duration of phonetic processing of the final speech sound of the word.

Together, these results confirm a key prediction of HDC: the dynamics of processing are increasingly sustained as the feature under consideration is high in the language hierarchy. We also observe that information at each level is encoded well into the processing of subsequent phonemes and words, leading to significant parallel processing, across and within levels of representation.

### 1.3. Hierarchical features are encoded in a dynamic neural code

We find that each linguistic feature can be decoded – and is thus represented – for a longer time window than its actual duration in natural speech.Here, we test the Hierarchical Dynamic Coding Hypothesis: a dynamic neural code allows successive phonemes, syllables and words to be maintained without representational overlap between sequential neighbours.

To test this, we implemented a temporal generalization analysis ^3,43^ (Figure 1). This method involves evaluating whether the topographic pattern learnt at time *t* generalizes to subsequent and preceding time-points (see Methods for details). If the representation is held within the same neural pattern over time, then the topographic pattern learnt at time *t* should generalize to time *t*+N, leading to a ‘square’ decoding matrix. By contrast, if the neural code evolves as a function of time, then the topographic pattern learnt at time *t* would not be the same at time *t*+N, even if the representation can also be decoded at t+N. In this scenario of a dynamic code, we thus expect to detect a ‘diagonal’ matrix.

Of primary interest are two parameters of this generalization matrix: (1) the window during which the representation can be decoded and (2) the window during which decoders tend to generalize.

We applied this analysis to each of our language features, and then averaged the generalization matrices over the six levels of interest (Figure 5) to estimate the similarity of spatial evolution across the hierarchy.

**Figure 5:**
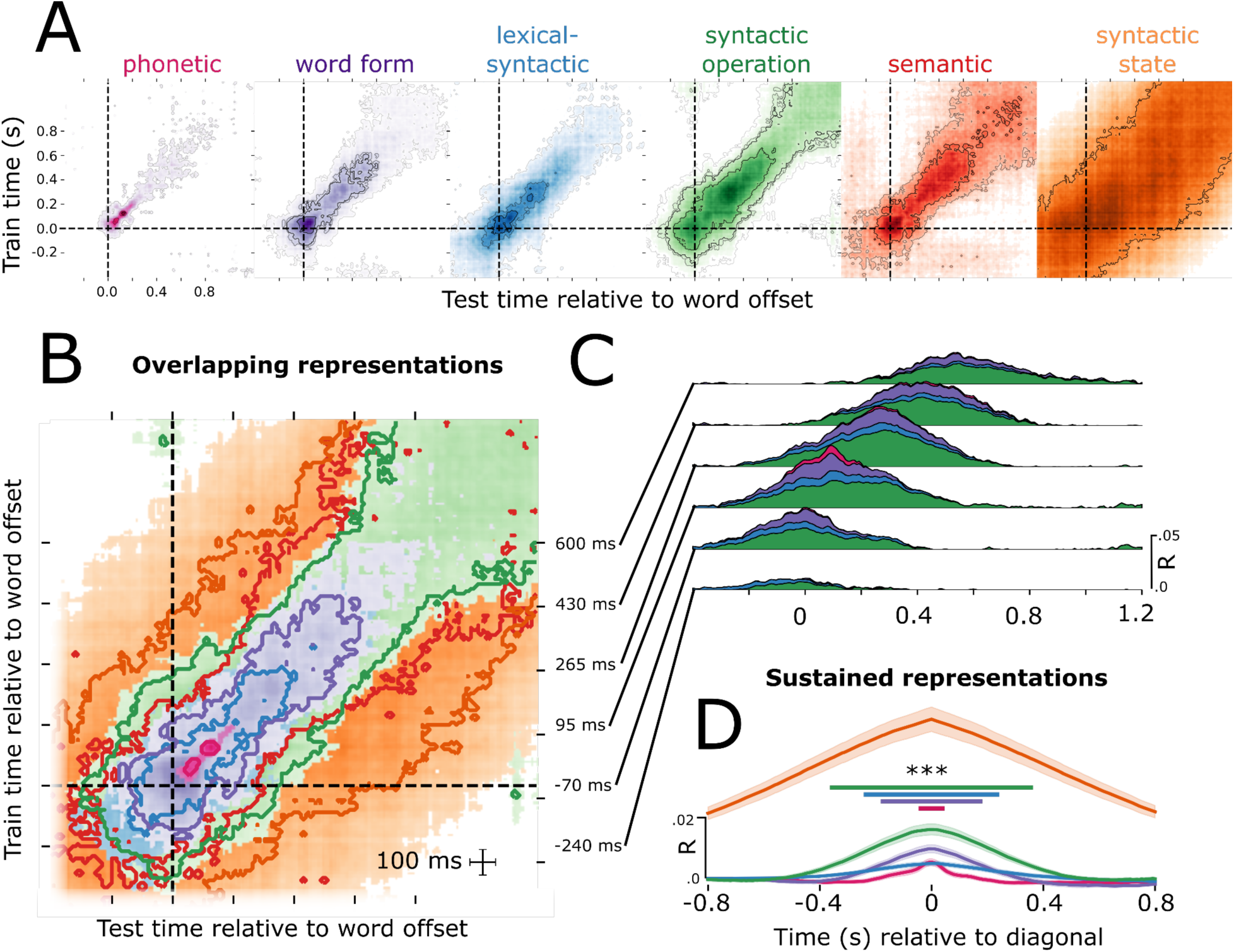
Evidence for Hierarchical Dynamic Coding. A: Temporal generalization analysis for each of the six linguistic levels of analysis. Each contour represents a significance threshold of p < .05, p < .01, p < .005, and p < .001. B: Same data as shown in A but for just the p < .01 threshold. C: Cumulative temporal generalization performance for the temporal decoders trained at different time-points relative to word offset, just for phonetic, word form, lexical-syntactic and syntactic operation. D: Data re-aligned relative to the diagonal of the temporal generalization matrix, showing the relationship between format maintenance and feature complexity, here just for phonetic, word form, lexical-syntactic, syntactic operation and syntactic state. ***: *p* < .001.

We found that all six feature families are processed using a dynamic neural code. The neural activity patterns associated with each linguistic features are only stable for a relatively short time period: phonetic duration=184 ms; sustain=64 ms; word form duration=752 ms; sustain=384 ms; lexical-syntactic duration=536ms; sustain=224 ms; syntactic operation duration=1392ms; sustain=720 ms; syntactic state duration=1250 ms; sustain=1600 ms). This means that all levels of representation across the hierarchy are supported by neural patterns that change over time.

Furthermore, the stability of a linguistic feature depends on its level in the language hierarchy: The lower-level phonetic features, which are defined over smaller linguistic units (phonemes), evolved significantly faster (average generalization time 64 ms), than lexical features (224 ms), and those, faster than syntactic features (730 ms). This led to a significant correlation between the location of the family in the hierarchy, and duration of information sustain (*r* = −0.89, *p* = .034) (Figure 5D). This finding suggests that while all levels of the hierarchy share a dynamic coding scheme, the speed with which information is routed to different neural patterns scales with unit duration and abstraction.

### 1.4 Simulating Hierarchical Dynamic Coding

Our results have revealed a number of potentially important components of the spatio-temporal dynamics of language encoding, and how they vary across the language hierarchy. In this final section, we perform a number of simulations to test the assumptions of the computational framework we are proposing.

First, we repeated our hierarchical analyses on the Mel spectrogram of the speech signal to test to what degree the hierarchy of language features are linearly encoded in the acoustics that enter the ear. We computed the power in 50 log-spaced frequency bands of the spoken stories, spanning from 1-5000 Hz (see Methods). From this spectral representation, we used Ridge regression to decode each of the 54 hierarchical language features described above, using the temporal generalization analyses described in section 1.3. We find that phonetic and word form features can be decoded from the Mel spectrogram better than chance, as confirmed with a random-shuffle permutation test (p < .001). We also found that Syntactic Operation could be decoded late in the epoch time-window, and Syntactic State could be decoded early in the epoch time-window (both p < .001). Upon further inspection, we identified that this is caused by systematic co-occurrence with onsets from silence and offsets into silence (see Supplementary Figure 10). Lexical-syntactic and the semantic vectors were not decodable from the Mel spectrogram at any latency in the epoch. Together this suggests that (i) lower level properties of speech are indeed linearly encoded in the acoustic input; (ii) seemingly higher-order syntactic features have acoustic correlates, linked to the beginnings and ends of sentences; (iii) lexical-syntactic and semantic features are not robustly encoded in the input (see Figure 6B, top row; Supplementary Figure 9). Overall this supports that our results are not a trivial reflection of the input, but rather reflect the outcome of an active neural process applied to that input.

**Figure 6:**
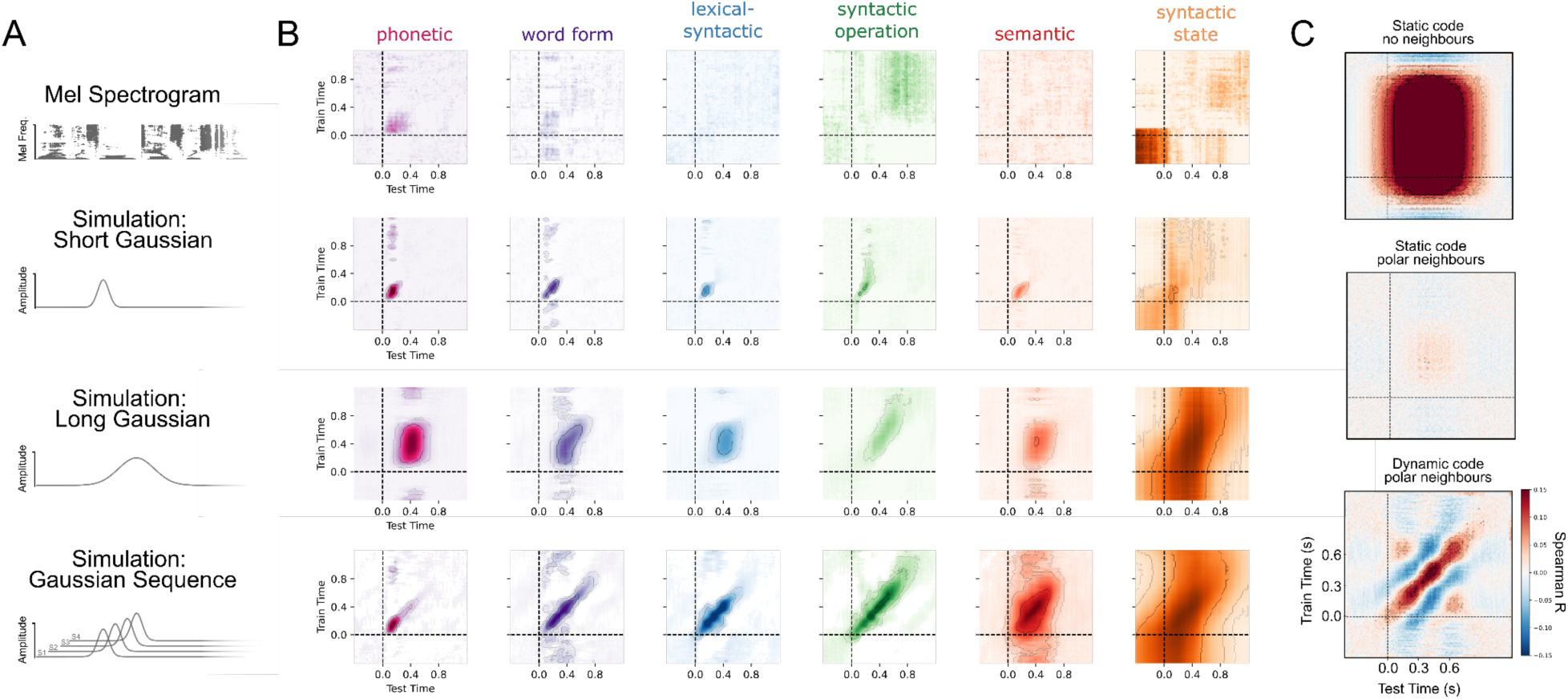
Simulating empirical results. A: We performed four simulations: on the Mel spectrogram of the speech; simulating a short gaussian response at linguistic feature onset; simulating a longer gaussian response at linguistic feature onset; simulating a sequence of gaussian responses, in line with the Hierarchical Dynamic Coding (HDC) hypothesis). B: Results of temporal generalization analysis when applied to each of the simulated datasets. C: Simulating static and dynamic neural codes, under conditions of sequence interference.

Second, we tested whether the dynamics we observe in Figure 4A and 5A could be a consequence of the dynamics of the language features in sparsity or autocorrelation. For example, language properties at the “top” of the hierarchy such as syntactic depth have a higher autocorrelation, given that the value of depth at the current word is likely to be correlated with subsequent and preceding words. We simulated MEG responses to the features in our stories, preserving all native feature dynamics. We performed three simulations: (i) spatially static short gaussian response; (ii) spatially static long gaussian response; (iii) spatially dynamic sequence of gaussian responses across sensors. For (i) we simulated each gaussian by selecting a peak time uniformly sampled from 100-200ms; an amplitude uniformly sampled from −1 to +1, and a width uniformly sampled from 20-50 ms. For (ii) we simulated each gaussian by selecting a peak time uniformly sampled from 350-500ms; an amplitude uniformly sampled from −1 to +1, and a width uniformly sampled from 100-150 ms. For (iii) we used the gaussian parameters from (i) but additionally added a cascade of gaussian responses, moving to distinct sensors each time. For each level of the hierarchy, we added an additional gaussian response to the sequence, each spaced 50 ms apart, thus simulating the hypothesised neural code of Hierarchical Dynamic Coding as outlined in Figure 1.

Our simulations (i) & (ii) reveal that for all features other than Syntactic State, the onset and duration of feature encoding directly reflected the onset and duration of the ground-truth neural response (e.g., see Supplementary Figure 11&12), and critically did not scale with the position of the level in the hierarchy. This means that the dynamics we observe empirically from our MEG decoding are crucially not merely a reflection of the dynamics of the stimulus features, but are the consequence of the spatio-temporal dynamics of the neural code applied to those features. In simulation (iii) we recapitulate our finding that a Hierarchical Dynamic Code gives rise to increased duration of encoding across each level of the hierarchy. We also find that only under the HDC simulation do we observe a “diagonal” generalization pattern, again providing evidence that the dynamics are not merely a consequence of the input.

The one exception to the above is the simulation of Syntactic State. We found that the duration of syntactic state encoded was much longer lived than the ground truth response vector. We attribute this to the extreme autocorrelation of this feature across word sequences. Consequently, we interpret the dynamics of this feature with caution, given that the prolonged encoding can also be recovered from a ground-truth transient gaussian response. As noted, Syntactic State is the only feature with this property.

### 1.5 Simulating Destructive Interference

That the neural code for shorter units evolved faster, and the neural code for longer units evolved slower, has the consequence that neighbouring units avoid representational overlap across the linguistic hierarchy. In this final analysis, we test whether this serves to avoid “destructive interference” whereby the features of neighbouring words would serve to cancel each other out, if the neural encoding of those words was shared. We define “interference” specifically as the case where two or more features of the input are encoded in neural activity at the same time, but their feature values are contrastive, thus leading to cancellation.

Here we test our hypothesis that a static and prolonged neural code would lead to catastrophic interference between neighbouring features. We simulated MEG responses as gaussian activation functions, with a peak response at 400 ms, amplitude of 1.5 femto-tesla, and response width equal to the average word duration in our stories (293 ms). We simulated responses of this static code using (i) exaggerated distance of 2 silent seconds between neighbouring words; (ii) actual distance between words from our story stimuli. To model maximal interference, we used a simulated feature vector that fluctuated between +1 and −1 at the onset of each word in the story. Finally, to simulate responses under the Hierarchical Dynamic Coding hypothesis, we encoded the maximally contrastive simulated feature in a sequence of gaussian responses that travel across space. Each gaussian in the sequence had a peak response at 400 ms, amplitude of 1.5 femto-tesla, and response width equal to the average word duration in our stories (293 ms). There were 3 gaussians in the sequence, occurring at 0 ms, 50 ms and 100 ms relative to feature onset.

We determined whether representational interference had occurred by decoding the ground-truth simulated vector back from each of the three simulated MEG responses. We applied temporal generalization analyses to our simulated MEG responses, and we found that when the silent spaces between neighbouring words were sufficiently exaggerated, the simulated feature vector could be accurately reconstructed. By contrast, when we used the actual rapid pace of words in our stories, as is representative of real speech, we found evidence for catastrophic interference, and the underlying feature vector could no longer be recovered. Finally, when the feature was encoded in a dynamic code that processed across space, the ground-truth contrastive vector was again recoverable (Figure 6C).

## Discussion

Speech comprehension hinges on a delicate balance between *maintaining* low-level elements – like words, long enough to integrate them into more complex units – like phrases^47^, while continuously *updating* each element in time with rapidly incoming information^34,41,42^. The neural implementation of this process must enable multiple features across the hierarchy, across an extended history, to be encoded simultaneously, while avoiding destructive interference between neighbouring inputs^48^.

We propose a new processing model that satisfies these constraints: Hierarchical Dynamic Coding (HDC). Our computational account, and associated neural description, challenges traditional models that posit a one-to-one mapping between a neural pattern and language feature^13,14^. Rather than a singular response latency^26,49^ at a given spatial location^9,50^, we find that language representations are encoded in a series of spatial patterns over time, and the full hierarchy is encoded largely in parallel. This provides an updated neuroscientific framework for interpreting the substantial body of behavioural research that demonstrates longevity of hierarchical representations^37–39^.

Given that the dynamic neural code traverses neural patterns in a systematic sequence, each neural pattern is “time-stamped” for absolute elapsed time since feature onset. This is similar to the rotary position embedding (RoPE) used in speech and language models^51,52^, whereby a continuously evolving projection helps the model to temporally locate, for example, the relative position of each word in a sentence, without the need for explicit ordinal position encoding^53^. While we do not leverage transformer models in the present study, it would be interesting to explicitly evaluate the similarity between the RoPE mechanism and the evolving dynamic code we report as part of HDC.

We also observe that this dynamic neural code evolves at a speed commensurate with the corresponding hierarchical level. In other words, the neural code shifts between neural patterns more slowly for high level representations than for low level representations. In addition, higher level features are maintained for significantly longer in the neural signal, extending into the processing of multiple words in the future. This neural overlap is much greater than previously appreciated ^2,9^, and may serve to build a sequence of sufficient length to build representations one level above, when operating in a bottom-up processing regime. Our simulation analyses confirm that the dynamical speed of evolution, and the duration of encoding, are not trivial consequences of the dynamics of the features themselves, but rather are the result of an active process that the brain applies to these neural representations. Anatomically, these results align with the finding that higher-level areas (e.g. fronto-parietal cortices) integrate speech representations over longer time periods than lower-level areas (e.g. Heschl’s gyrus) ^29,36,54–56^.

Notably, the two syntactic levels–syntactic state and syntactic operation–revealed significantly different dynamics. The syntactic operation level, which includes features such as number of opening and closing nodes, evolved more quickly and was sustained for less time than syntactic state. Our simulation analyses help to clarify this difference: Even when we only simulate a single gaussian response to the syntactic state feature (**Figure 6**), the decoding dynamics out-live the ground truth neural generators. This is due to the extremely high autocorrelation of the syntactic state feature as compared to the other features. In addition, syntactic state was significantly decodable from the Mel spectrogram, which we show is due to co-variance with sentence offsets. As a consequence, our acoustic control analysis, and simulation results, show that the dynamics of syntactic state are partially driven by the autocorrelation of the features at that level, and should be interpreted with caution. None of the other feature dynamics were explainable in this way–suggesting that the dynamics at all other hierarchical levels are indeed due to dynamics of the given neural process.

A major computational advantage of adjusting evolution speed with hierarchical level is the avoidance of destructive interference between neighbouring elements of the speech input. If two or more features of the input are encoded in the same neural activity pattern, and their feature values are contrastive, we show in simulation that this prevents robust recovery of the underlying sequence representation. Given that phoneme sequences occur in more rapid succession than word or phrasal sequences, it is necessary for the dynamic code of phonemes to evolve more quickly to avoid representational overlap. When we simulate speech with exaggerated silences between features, static code destructive interference is resolved; however, the speech slow-down is suboptimal from an efficient communication standpoint^57^, and would require information to be maintained for significantly longer periods. Thus, HDC provides an explicit account of how this dynamic code allows speech to unfold rapidly while serving to avoid destructive interference between neighboring inputs.

We observe that all levels of linguistic encoding, from sound to meaning, are supported by a dynamic neural code. This raises the question of whether this dynamic code may be observable for other sequential stimuli, and in other sensory modalities. Previous work that focuses on just one level of representation has shown that novel visual stimuli, either presented in isolation^58^ or presented in quick succession^46^ as well as non-speech frequency-modulated tones ^59^ also elicit a dynamic “cascade” of activity. This suggests that the dynamic code is not only recruited for highly trained stimuli, or only for auditory stimuli, but rather this is a “canonical” process that is applied across multiple processing domains. Previous work^59^ associated different points in the response to auditory tones with sensitivity to different aspects of the experimental design: sensory ambiguity and contextual prediction. This suggests that the information that is encoded in the evolving neural pattern may also be changing over time – but, all “versions” of information are sufficiently correlated with our feature probes to provide successful readout throughout the response timecourse. Key questions to address in future work are whether, and in what ways, the information at each hierarchical level is changing over time. In addition, intracranial data would provide precise insight into *how* the spatial pattern evolves: how “far” does the neural code move; is it primarily within a brain region, or does information move across brain regions? Answers to both questions would help build a detailed understanding of the formats of information that are available to downstream areas at different latencies in processing, as well as the neural computations in place that modify the representational format through this processing pipeline.

Previous research investigating the dynamics of language feature processing using scalp EEG have studied many of the same levels of representation we explore here, by manipulating the expectation of one hierarchical dimension at a time, and observing corresponding temporal fluctuations in activity strength^60^. Studies report that expectancy of lower order phonetic features^31^ lead to responses earlier than manipulations of predictably higher levels, including semantics and syntax^27,28,61,62^. Our findings contrast with these prior results, showing that the onset of higher level feature encoding actually precedes the onset of lower feature encoding. We reconcile these findings by noting an important difference in our approach: we are decoding the *value* of the feature directly (e.g., “Noun”) rather than the surprisal of that feature (e.g., log probability of Noun). Studies exploring EEG responses during continuous speech have found that surprisal across the hierarchy is aligned with more traditional EEG studies–that is, surprisal of phonetics precedes surprisal of semantic and syntax–and they do not observe the “reverse” hierarchical encoding pattern we report here. Another difference is that we locked our analysis to word offset rather than word onset, because it consistently yielded more robust decoding performance; however, when we lock our analysis to word onset, we find a qualitatively similar pattern. This suggests that it is our choice of feature set, and not our use of a naturalistic paradigm, or choice of event-locking, explains the difference in dynamics. Future studies should include both direct feature representations (as used in the current study) and surprisal over those feature representations (as is more typical in the EEG literature), to reconcile the relative time courses and extent of neural overlap.

The early feature encoding we observe is interesting, as it could be an indication of the brain’s *prediction* about the upcoming feature, which is later evaluated relative to the actual feature outcome– leading to the surprisal response^63,64^. This is in line with recent EEG studies investigating “pre-activation” of expected feature outcomes, using highly constraining sentences^65,66^. This account forms the hypothesis that when the listener is hearing the beginning of the story, the onset of feature encoding must proceed in the more traditional bottom-up order, given that there is minimal context to leverage for predictions. As the story unfolds, the brain can increasingly form predictions about upcoming inputs, thus leading to increasingly earlier feature encoding. Future studies should test whether this account bears out–that the direction and steepness of onset decoding directly scale with the predictability of upcoming inputs.

That increasingly higher levels have increasingly earlier onset leads to the question of whether higher order structures actually serve to feed their predictions to lower levels, in order to guide and bias how lower level representations are interpreted^67^. This is consistent with the “Good Enough” and “Syntax First” models of language processing^68–71^, where higher order structures are used to guide comprehension based on what is coherent in context. This “reverse” architecture of speech processing presents three computational advantages^72,73^. First, because higher-level language features are abstracted away from the sensory signal, comprehension is more robust to auditory noise and ambiguity ^74,75^. Second, ambiguity at one level of representation may be resolvable by integrating information from other levels^76 6,77,78,79^. Finally, it potentially speeds up processing by initiating high-level computations early during comprehension, rather than waiting for them to be formed compositionally in purely bottom-up fashion^68,69,71^.

Overall, our results offer an updated computational account, and associated neural description, of how the brain maintains and updates the continuously unfolding hierarchical representations of spoken language. We track a comprehensive hierarchy of features, ranging from phonemes to syntactic trees, and find evidence for a canonical dynamic code, which adapts its processing speed as a function of level in the language hierarchy. Hierarchical Dynamic Coding (HDC) elegantly balances the preservation of information over time with minimizing overlap between consecutive language elements. This system provides a clear view of how the brain may organize and interpret rapidly unfolding speech in real time, linking linguistic theories with their neurological foundations.

## Acknowledgements

We thank Graham Flick for help with data collection. A big thanks to Kara Federmeier, Florencia Assaneo,

Joan Opella, Arianna Zuanazzi, Suzanne Dikker and Jill Kries for feedback on a previous version of the manuscript.

## Author contributions

LG: conceptualisation; methodology; software; validation; formal analysis; investigation; data curation; writing - original draft preparation and review and editing; visualization. JRK: conceptualisation; methodology; software; supervision. AM: conceptualisation; writing - review and editing; supervision; funding acquisition. DP: conceptualisation; writing - review and editing; supervision; funding acquisition.

## Data and materials availability

Preprocessed data have been publicly released on the Open Science Framework (https://osf.io/ag3kj/, ^80^).

## Methods

We note that we are analysing a naturalistic dataset of participants listening to short stories, which was also analysed in a previous study from these authors: Gwilliams et al., 2022, *Nature Communications*. In the previous study, we focused our analysis purely on the acoustic and phonetic levels of processing, and analysed the responses to 50,518 phonemes per participant. Here, we are focusing our analyses on the 13,798 words and analysing responses relative to a comprehensive *hierarchy* of language representation–significantly moving beyond the level of acoustic-phonetics. We note that all of the data preprocessing steps we outline below are the same as described in our prior paper. But the features we explore, the analyses we apply, and the simulations we run, are unique to the current paper.

### 3.0. Definition of terms

- **Feature**: a property of language: e.g., “fricative” phonetic feature; word frequency, etc.
- **Level**: a group of features at the same degree of hierarchical position: e.g. phonetic, word form, lexical-syntactic, syntactic operation, semantic, syntactic state
- **Representation**: neural encoding scheme of a feature
- **Dynamic coding**: when neural representations are instantiated by distinct neural patterns over time.
- **Hierarchical Dynamic Coding**: when different levels follow a dynamic coding scheme

### 3.1. Participants

Twenty-one native English participants were recruited from the NYU Abu Dhabi community (13 female; age: M=24.8, SD=6.4). All provided their informed consent and were compensated for their time. Participants reported having normal hearing and no history of neurological disorders. Each subject participated in the experiment twice. Time between sessions ranged from 1 day to 2 months. All participants gave their informed consent, and the experiment was approved by the local IRB committee of NYU Abu Dhabi.

### 3.2. Stimulus development

Four fictional stories were selected from the Open American National Corpus ^81^: Cable spool boy (about two bothers playing in the woods); LW1 (sci-fi story about an alien spaceship trying to find home); Black willow (about an author struggling with writer’s block); Easy money (about two old friends using magic to make money).

Stimuli were annotated for phoneme boundaries and labels using the ‘gentle aligner’ from the Python module *lowerquality*. Some prior testing provided better results than the Penn Forced Aligner ^82^. To verify that the forced alignment did not have a systematic bias, we checked the MEG decoding of phonetic features for each sound file separately. In some cases, the aligner failed to find an alignment; we removed all such words from all analyses presented here.

Each of the stories were synthesised using the Mac OSX text-to-speech application. Three synthetic voices were used (Ava, Samantha, Allison). Voices changed every 5-20 sentences. The speech rate of the voices ranged from 145-205 words per minute, which also changed every 5-20 sentences. The silence between sentences randomly varied between 0-1000 ms.

### 3.3. Procedure

Before the experiment proper, the participant was exposed to 20 seconds of each speaker explaining the structure of the experiment. This was designed to help the participants attune to the synthetic voices. The order of stories was fully crossed using a Latin-square design. Participants heard the stories in the same order during both the first and second sessions. This was in order to make direct comparisons between the first and second sessions.

Participants answered a two-choice question on the story content every ∼3 minutes. For example, one of the questions was “what was the location of the bank that they robbed”? The purpose of the questions was to keep participants attentive and to have a formal measure of engagement. All participants performed this task at ceiling, with an accuracy of 98%. Participants responded with a button press. Stimuli were presented binaurally to participants though tube earphones (Aero Technologies), at a mean level of 70 dB SPL. The stories ranged from 8-25 minutes, with a total running time of ∼1 hour.

The stimuli used for this study are detailed in Gwilliams et al., (2023) *Scientific Data*^80^. The stimuli of text, sound and timing are available on https://osf.io/ag3kj/.

#### Stories

Each participant listened to four fictional stories, over the course of two ∼1h-long MEG sessions, with the exception of 5 subjects who only underwent 1 session. The stories were played in different orders across participants. These stories were originally selected because they had been annotated for their syntactic structures^81^. The corresponding text files can be found in stimuli/text/*.txt

#### Word lists and pseudo-words

To potentially investigate MEG responses to words independently of their narrative context, the text of these stories have been supplemented with word lists. It is common to use “word lists” as a baseline condition in labs using fMRI to study language processing (e.g., for a review see ^83^. Specifically, a random word list consisting of the unique content words (nouns, proper nouns, verbs, adverbs and adjectives) selected from the preceding text segment was added in a random order. In addition, a small fraction (<1%) of non-words were inserted into the natural sentences of the stories. Again, the reason here is because non-words are also often used as a baseline comparison condition in neurolinguistics. The corresponding text files can be found in stimuli/text_with_wordlist/*.txt.

#### Importantly, the brain responses to these word lists and to these pseudo words are fully discarded from the present study. This had no negative bearing on the accuracy of the timestamps

##### Audio synthesis

Each of these stories was synthesized with Mac OS Mojave © version 10.14 text-to-speech. Voices (n=3 female) and speech rates (145 - 205 words per minute) varied every 5-20 sentences. The inter-sentence interval randomly varied between 0 and 1,000 ms. Both speech rate and inter-sentence intervals were sampled from a uniform distribution. Each ‘text_with_wordlist’ files was divided into ∼3 min sound files, which can be found in stimuli/audio/*.wav.

##### Forced Alignment

The timing of words and phonemes were inferred from the forced-alignment between the wav and text files, using the ‘gentle aligner’ from the Python module lowerquality (https://github.com/lowerquality/gentle). We discard, from subsequent analyses, the words that did not get a forced alignment through this procedure. Because the aligner uses a context window to determine alignment, there were some moments where uncertainty in the alignment of a single word resulted in full sentences being absent from the alignment^84^. Analysis of the Mel spectrogram and of the phonetic decoding led to better results when using gentle than when using the Penn Forced Aligner. The timing of each word and phoneme can be found in the events.tsv of each individual recording session.

##### Verification

To verify that the forced alignment did not have a systematic bias, we checked the MEG decoding of phonetic features for each sound file separately.

### 3.4. MEG acquisition

Marker coils were placed at the same five positions to localise each participant’s skull relative to the sensors. These marker measurements were recorded just before and after the experiment in order to track the degree of movement during the recording.

MEG data were recorded continuously using a 208 channel axial gradiometer system (Kanazawa Institute of Technology, Kanazawa, Japan), with a sampling rate of 1000 Hz and applying an online low-pass filter of 200 Hz.

### 3.5. Preprocessing MEG

The raw MEG data were noise reduced using the Continuously Adjusted Least Squares Method (CALM: ^85^, with MEG160 software (Yokohawa Electric Corporation and Eagle Technology Corporation, Tokyo, Japan).

The data were bandpass-filtered between 0.1 and 50 Hz using MNE-Python’s default parameters with firwin design [50] and downsampled to 250 Hz. We used MNE-Python version 1.3.0.

We segmented the data into epochs, from 400 ms pre-word onset to 1200 ms post-word onset. This modifies the data structure from being of [sensors x continuous_time] into [sensors x 1600ms x words]. No baseline correction was applied. This epoching step facilitates the decoding analysis method. Because we fit a Ridge regression decoding model at each lag separately relative to the boundary between words. Epoching the data relative to word offset (in the Main analysis) and relative to word onset (in the Supplementary analysis) makes the analysis procedure convenient to implement algorithmically, because we can fit the Ridge regression on each time lag in parallel.

### 3.6. Effects of acoustic features

In order to promote our decoding algorithm to identify neural patterns that correspond to language encoding per se, rather than low-level acoustic fluctuations in amplitude and pitch, we decided to regress out these features from the MEG data before applying our decoding method. When comparing the results to the non-regressed data, we found that this step did not make a difference to the interpretations of our results; however, given that it was a part of our initial analysis plan, we decided to keep this processing step to avoid introducing additional experimental degrees of freedom.

We used a temporal receptive field (TRF) model to regress from the raw MEG data responses that were sensitive to fluctuations in the pitch and envelope of the acoustic speech signal. We used the *ReceptiveField* function from MNE-Python ^86^, using ridge regression as the estimator and laplacian regularization. We tested ten lambda regularization parameters, log-spaced between 1^−6^ and 1^+6^, and picked the model with the highest predictive performance averaged across sensors. MEG sensor activity at each ms were modeled using the preceding 200 ms of envelope and pitch estimates. Both the acoustic and MEG signals were demeaned and scaled to have unit variance before fitting the model. MEG acoustic-based predictions were then transformed back into original MEG units before regressing out of the true MEG signals. This process, including fitting hyper-parameters, was applied for each story recording and for each subject separately, across 3 folds. This yields a de-confounded MEG dataset on which to continue our analysis.

### 3.7. Modeled features

We investigated whether single-trial sensor responses varied as a function of 54 features. Features spanned different levels of the linguistic hierarchy and included both binary and continuous variables.

#### 3.7.1. Phonetic

Phonetic features were derived from the multi-value feature system reported in ^87^. Note that this feature system is sparse relative to the full set of distinctive features that can be identified in English; however, it serves as a reasonable approximation of the phonemic inventory for our purposes.

##### Voicing

This refers to whether the vocal cords vibrate during production. For example, this is the difference between *b* versus *p* and *z* versus *s*.

##### Manner of articulation

Manner refers to the way by which air is allowed to pass through the articulators during production. Here we tested five manner features: fricative, nasal, plosive, approximant, and vowel.

##### Place of articulation

Place refers to where the articulators (teeth, tongue, lips) are positioned during production. For vowels, this consists of: central vowel, low vowel, mid vowel, high vowel. For consonants, this consists of: coronal, glottal, labial and velar.

| Feature Category | Description | Feature Values | Examples |
| --- | --- | --- | --- |
| Voicing | Whether the vocal cords vibrate during production. | **Voiced** (vocal cords vibrate) | **Voiced:** b, d, g, z |
|  |  | **Voiceless** (no vibration) | **Voiceless:** p, t, k, s |
| Manner of Articulation | How air passes through the articulators during speech. | **Fricative** (continuous airflow with friction) | **Fricative:** f, s, v, z |
|  |  | **Nasal** (air passes through the nose) | **Nasal:** m, n |
|  |  | **Plosive** (airflow is stopped, then released) | **Plosive:** p, t, b, d |
|  |  | **Approximant** (articulators come close, but no friction) | **Approximant:** w, r, l, j |
|  |  | **Vowel** (open vocal tract, no significant constriction) | **Vowel:** a, e, i, o, u |
| Place of Articulation (Vowels) | Where the tongue is positioned in the mouth. | **Central** (tongue in the middle) | **Central:** ə (schwa) |
|  |  | **Low** (tongue positioned low) | **Low:** æ (as in *cat*) |
|  |  | **Mid** (tongue positioned mid-height) | **Mid:** e (as in *bet*), o (as in *bore*) |
|  |  | **High** (tongue positioned high) | **High:** i (as in *see*), u (as in *blue*) |
| Place of Articulation (Consonants) | Where the articulators (tongue, teeth, lips) shape the sound. | **Coronal** (tongue tip/blade) | **Coronal:** t, s, d, z |
|  |  | **Glottal** (vocal cords) | **Glottal:** h |
|  |  | **Labial** (lips) | **Labial:** p, b, m |
|  |  | **Velar** (back of tongue) | **Velar:** k, g, ŋ (as in *sing*) |

#### 3.7.2. Word Form

To compute word form features, we used the English Lexicon Project (ELP) ^88^, which is a comprehensive database of English words that provides detailed phonological, morphological, and frequency-based metrics, which includes a corpus of spoken English. The ELP corpus includes a large collection of words from spoken and written English, allowing for a systematic quantification of various linguistic properties.

One key set of features includes the number of phonemes, syllables, and morphemes in each word. These measures provide a structural breakdown of the word into its constituent units, capturing its complexity at different linguistic levels. Phonemes represent the smallest units of sound that distinguish meaning, syllables structure the pronunciation and rhythm of a word, and morphemes reflect its smallest meaningful components, including roots and affixes. Words with more phonemes or syllables may be more challenging to articulate and process, while those with more morphemes may carry greater semantic complexity due to their morphological composition. These counts allow for a finer analysis of how word structure influences cognitive and linguistic processing.

A related measure is the number of phonemes within a syllable, which reflects the internal complexity of a word’s syllabic structure. Some syllables contain only a vowel (e.g., “a”), while others have multiple consonants clustered together (e.g., “strength”). This measure provides insight into the phonological density of words, which can impact pronunciation difficulty and processing time in both spoken and written language.

In addition to structural features, we also considered word frequency, which was derived from the subtitles corpus of American English within the ELP. Rather than using raw frequency counts, we applied a log transformation to normalize the distribution, as word frequency follows a highly skewed pattern— some words appear extremely often (e.g., the, is), while others are much rarer (e.g., serendipity, quixotic). Word frequency is a crucial predictor in psycholinguistics, as high-frequency words tend to be recognized and retrieved more quickly and accurately compared to low-frequency words. This effect is well-documented in lexical decision tasks, reading studies, and spoken word recognition research.

Finally, we incorporated phonological neighborhood density, a measure of lexical competition. This metric captures how many other words can be formed by changing a single phoneme in the target word. For example, “cat” has many phonological neighbors (bat, mat, cut), making it part of a dense neighborhood, while “orange” has very few close phonological relatives and is in a sparse neighborhood. Words in dense phonological neighborhoods tend to experience more competition during retrieval, potentially slowing recognition and increasing processing difficulty. This effect has been widely studied in models of spoken word recognition and lexical access, as it influences both comprehension and speech production.

Together, these features provide a detailed linguistic profile of words, integrating phonological, morphological, and frequency-based dimensions. Their inclusion is essential for studying how different properties of words influence cognitive processing, whether in psycholinguistic experiments, computational models, or applied linguistic research.

#### 3.7.3. Lexical-syntactic

Our lexical-syntactic level contains part of speech labels for every word. We derived these labels from the syntactic parse of the stories. This means that each word was assigned a syntactic category based on its role within the sentence structure, following standard linguistic parsing techniques.

To facilitate analysis, these word class labels were dummy coded relative to a set of 11 distinct word-class categories. Dummy coding is a common technique in statistical modeling and machine learning, where categorical variables are represented as binary indicators (0 or 1) for each possible category. This allows for flexible and interpretable numerical representation of categorical linguistic features.

The 11 word-class categories included in our coding scheme are:

- Adjective – Descriptive words that modify nouns (e.g., blue, tall).
- Coordinating conjunction – Words that link clauses, phrases, or words of equal grammatical rank (e.g., and, or).
- Determiner – Words that introduce and specify nouns (e.g., the, a).
- Noun – Words representing people, places, things, or ideas (e.g., house, girl).
- Pronoun – Words that replace nouns to avoid repetition (e.g., she, they).
- Preposition – Words that indicate relationships between elements in a sentence, often expressing direction, place, or time (e.g., under, on).
- Adverb – Words that modify verbs, adjectives, or other adverbs, often describing manner, time, or degree (e.g., slowly, fast).
- Verbal preposition – A specific type of preposition that functions as part of verb constructions (e.g., to, as in want to go).
- Verb – Action words or states of being (e.g., run, jump).
- WH-Word – Question words used to form interrogative and relative clauses (e.g., where, who).
- Existential there – The use of there to indicate the existence of something (e.g., there is a book on the table).

#### 3.7.4. Syntactic operation

These features are derived from the penn-treebank syntactic parse of the stories. For example, the sentence “The cat sat on the mat” has the following Penn Treebank representation:

~~~
(S (NP (DT The) (NN cat)) (VP (VBD sat) (PP (IN on) (NP (DT the) (NN mat)))))
~~~

The number of closing nodes in a syntactic parse tree refers to the number of times a non-terminal node in the tree closes (i.e., when a constituent or subtree completes). This can be interpreted as the number of times a right parenthesis “)” appears in a Penn Treebank-style bracketed representation. Since each “)” represents the closure of a constituent, to extract the number of closing nodes from a Penn Treebank-style parse tree, we simply count the closing parentheses.

Similarly, the number of opening nodes in a syntactic parse tree refers to the number of times a new non-terminal node begins in a Penn Treebank-style representation. This corresponds to the number of times an opening parenthesis “(” appears in the bracketed structure. Again, to compute the number of opening nodes, we simply count the number of closing parentheses.

Note that these features are intended to be general constructs in Linguistics, and not tied to a specific theory.

#### 3.7.5. Syntactic state

These features are also derived from the Penn-Treebank syntactic parse of the stories.

We compute tree depth as the longest path from the root node to any leaf node. In a Penn Treebank-style parse tree, this corresponds to the maximum level of nested opening parentheses “(” at any point in the string. The current depth at a given word is provided by subtracting the count of closed parentheses “)” from the number of open parentheses “(”.

For example, below is the depth count in square brackets for each word in the sentence.

~~~
(S
 (NP (DT The[2]) (NN cat[2]))
 (VP (VBD sat[2])
  (PP (IN on[3])
   (NP (DT the[4]) (NN mat[4]))
  )
 )
)
~~~

The number of open nodes at a particular word in a Penn Treebank-style parse tree corresponds to how many opening parentheses “(” have been encountered without a matching closing parentheses “)” at that point in the string. In other words, it represents how many syntactic constituents are still “open” when the word appears in the tree.

| Step | Action | Open Nodes | Word Produced |
| --- | --- | --- | --- |
| (S | 1 | 1 | - |
| (NP | 1 | 2 | - |
| (DT | 1 | 3 | - |
| The | (Word) | 3 | The |
| ) | -1 | 2 | - |
| (NN | 1 | 3 | - |
| cat | (Word) | 3 | cat |
| ) | -1 | 2 | - |
| ) | -1 | 1 | - |
| (VP | 1 | 2 | - |
| (VBD | 1 | 3 | - |
| sat | (Word) | 3 | sat |
| ) | -1 | 2 | - |
| (PP | 1 | 3 | - |
| (IN | 1 | 4 | - |
| on | (Word) | 4 | on |
| ) | -1 | 3 | - |
| (NP | 1 | 4 | - |
| (DT | 1 | 5 | - |
| the | (Word) | 5 | the |
| ) | -1 | 4 | - |
| (NN | 1 | 5 | - |
| mat | (Word) | 5 | mat |
| ) | -1 | 4 | - |
| ) | -1 | 3 | - |
| ) | -1 | 2 | - |
| ) | -1 | 1 | - |
| ) | -1 | 0 | - (End) |

#### 3.7.6. Semantic vector

We obtained 50-dimensional word embedding GloVe vectors (references 61,62) for each word in our dataset. These embeddings represent words as points in a high-dimensional space, capturing semantic and syntactic relationships based on co-occurrence patterns in large text corpora. The GloVe vectors are trained using a symmetric context window of size 10, and therefore they capture meaning beyond the single lexical item.

To reduce the dimensionality of these embeddings while preserving as much meaningful variation as possible, we applied Principal Component Analysis (PCA). PCA is a statistical technique that transforms the original high-dimensional data into a lower-dimensional space by identifying orthogonal directions (principal components) that capture the most variance in the data.

As a result of applying PCA, we extracted the top ten principal components. These components are ordered according to the amount of variance they explain in the original 50-dimensional GloVe embeddings. The first principal component captures the largest proportion of variance, followed by the second, and so on. This organization ensures that the most critical dimensions of variation in the word embeddings are retained, while less significant noise and redundancy are minimized.

By leveraging these ten principal components, we obtain a lower-dimensional representation of word meanings that retains the most important semantic distinctions while reducing computational complexity and mitigating redundancy in the original embedding space.

### 3.8. Back-to-back regression decoding

We fit a back-to-back regression algorithm ^89^, which allows us to decode multiple features from the MEG data while also controlling for their co-variation. The resulting model coefficients represent how robustly a linguistic feature is encoded in neural responses, above and beyond the variance accounted for by the other linguistic features.

For the neural decoding, the input features were the magnitude of activity at each of the 208 MEG sensors. This approach allows us to decode from multiple, potentially overlapping, neural representations, without relying on gross modulations in activation strength ^89,90^.

Because some of the features in our analysis are correlated with one another, we need to jointly evaluate the accuracy of each decoding model relative to its performance in predicting all modeled features, not just the target feature of interest. This is because, if fitting each feature independently, we will not be able to dissociate the decoding of feature *f* from the decoding of the correlated feature *f*^. The necessity to use decoding over encoding models here, though (which, do not suffer so harshly from the problem of co-variance in the stimulus space) is one of signal to noise: we expect any signal related to linguistic processes to be contained in low-amplitude responses that are distributed over multiple sensors. Our chances of uncovering reliable responses to these features is boosted by using multivariate models. To overcome the issue of covariance, but still to capitalize on the advantages of decoding approaches, we implement a back-to-back ridge regression model [30]. This involves a two stage process. First, a ridge regression model was fit on a random half of the data, at a single time-point (steps 1-6 in Figure 7). The mapping was learnt between the multivariate input (activity across sensors) and the univariate stimulus feature (one of the 54 features described above). All decoders were provided with data normalized by the mean and standard deviation in the training set:

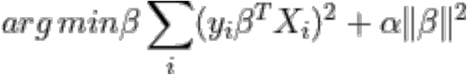

where *y_i_* ∈ {±1} is the feature to be decoded at trial *i* and *X_i_* is the multivariate neural measure. The l2 regularization parameter α was also fit, testing 20 log-spaced values from 1^−5^ to 1^5^. This was implemented using the *RidgeCV* function in *scikit-learn* ^91^.

Then, we use the other half of the acoustic or neural responses to generate a prediction for each of the 31 features corresponding to the test set. However, because the predictions are correlated, we need to jointly-evaluate the accuracy of decoding each feature, to take into account the variance explained by correlated non-target features. To do this, we fit another ridge regression model (steps 6-8 in Figure 7), this time learning the beta coefficients that map the matrix of *true* feature values to *predicted* feature values:

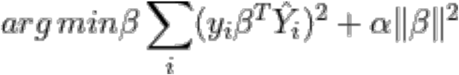

where *y_i_* ∈ {±1} is the ground truth of a particular stimulus feature at trial *i* and *Y*^*_i_* is the prediction for all stimulus features. A new regularization parameter α was learnt for this stage. By including all stimulus features in the model, this accounts for the correlation between the feature of interest and the other features. From this, we use the beta-coefficients that maps the true stimulus feature to the predicted stimulus feature. Beta coefficients serve as our metric of decoding performance: if a stimulus features is not encoded in neural responses (the null hypothesis) then there will be no meaningful mapping between the true feature *y* and the model prediction *y*^. Thus, the beta coefficient will be zero – equivalent to chance performance. If, however, a feature *is* encoded in neural activity (the alternative hypothesis), we should uncover a significant relationship between *y* and *y*^, thus yielding an above-zero beta coefficient.

The train/test split was performed over 100 folds, and the beta-coefficients were averaged across folds. This circumvents the issue of unstable coefficients when modeling correlated variables. These steps were applied to each subject independently.

Each of these steps are depicted graphically below in Figure 7.

**Figure 7:**
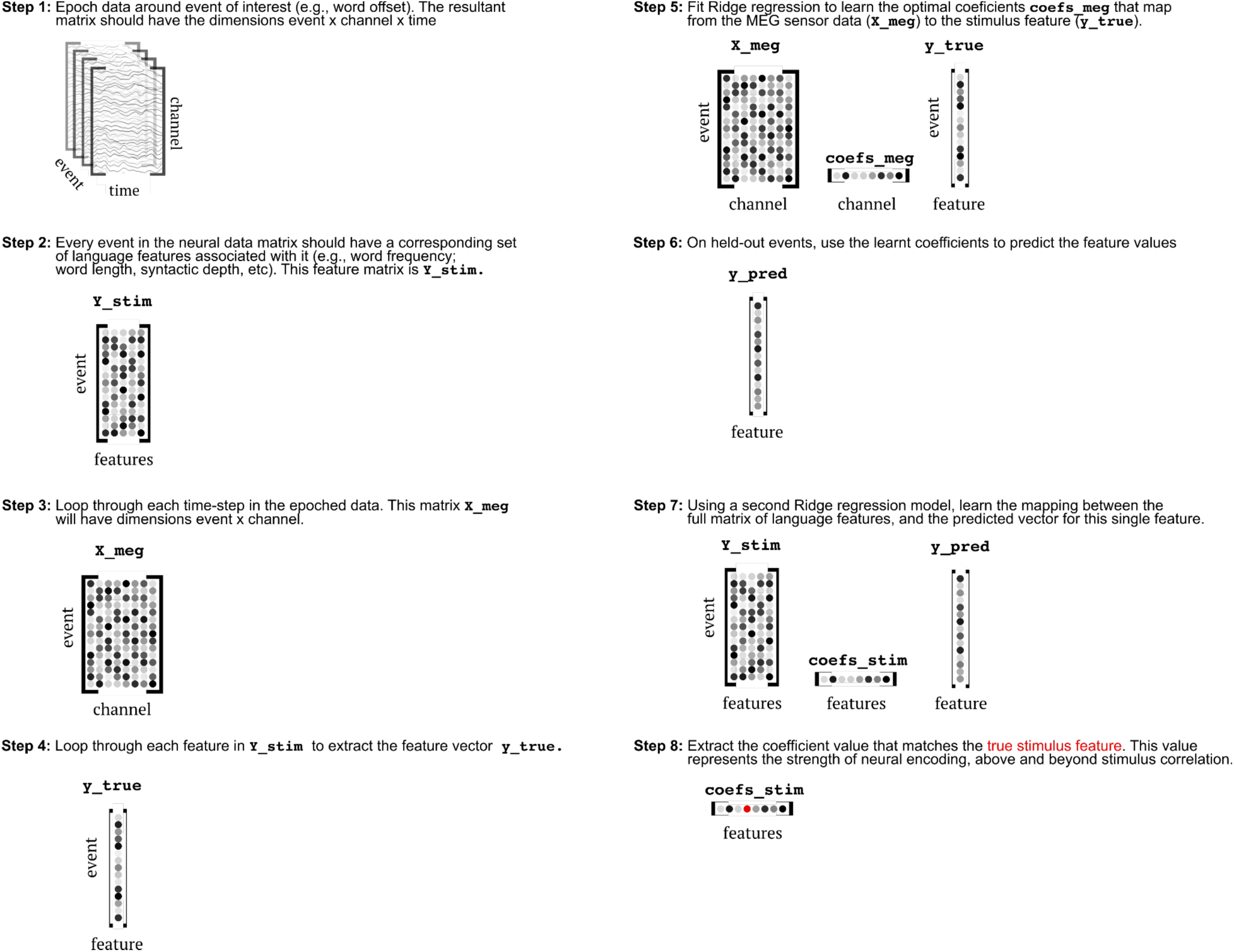
Steps of back-to-back regression analysis method.

### 3.9. Temporal generalization decoding

Temporal generalization (TG) consists of testing whether a temporal decoder fit on a training set at time *t* can decode a testing set at time *t*^′ 43^. This means that rather than evaluating decoding accuracy just at the time sample that the model was trained on, we evaluate its accuracy across all possible train/testing time combinations.

TG can be summarized with a square training time × testing time decoding matrix. To quantify the stability of neural representations, we measured the duration of above-chance generalization of each temporal decoder. To quantify the dynamics of neural representations, we compared the mean duration of above-chance generalization across temporal decoders to the duration of above-chance temporal decoding (i.e. the diagonal of the matrix versus its rows). These two metrics were assessed within each subject and tested with second-level statistics across subjects.

### 3.10. Comparing decoding performance between trial subsets

To evaluate whether the processing of syntax built over time, we subset our analysis by evaluating decoding performance at different word positions in the sentence. We add a modification to our train/test cross-validation loop. The data are trained on the entire training set (i.e. the same number of trials as the ‘typical analysis’), and the test set is grouped into the different levels of interest. We evaluate model performance separately on each split of the test data, which yields a time-course or generalization matrix for each group of trials that we evaluate on: in this case, each word position in the sentence.

### 3.11. Group statistics

To evaluate whether decoding performance is better than chance, we perform second-order statistics. This involves testing whether the distribution of beta coefficients across subjects significantly differs from chance (zero) across time using a one-sample permutation cluster test with default parameters specified in the MNE-Python package ^86^.

### 3.12. Simulation analyses

We simulated MEG responses to the features in our stories, using gaussian responses. The gaussian we formed by selecting a peak time uniformly sampled from a given time-window, an amplitude uniformly sampled from −1 to +1, and a width uniformly sampled from a given time window. For details on the peak time and width selected in each analysis, see the corresponding parameters in the results section.

To simulate a “static” response, we assigned the gaussian response encoding a given language feature to one MEG sensor. For a “dynamic” response, we encoded a given language feature in a sequence of gaussian responses, each spaced 50 ms apart.

To simulate the consequence of “destructive interference” under our different coding schemes, we simulated MEG responses as gaussian activation functions, with a peak response at 400 ms, amplitude of 1.5 femto-tesla, and response width equal to the average word duration in our stories (293 ms). We simulated responses of this static code using (i) exaggerated distance of 2 silent seconds between neighbouring words; (ii) actual distance between words from our story stimuli. To model maximal interference, we used a simulated feature vector that fluctuated between +1 and −1 at the onset of each word in the story. Finally, to simulate responses under the Hierarchical Dynamic Coding hypothesis, we encoded the maximally contrastive simulated feature in a sequence of gaussian responses that travel across space. Each gaussian in the sequence had a peak response at 400 ms, amplitude of 1.5 femto-tesla, and response width equal to the average word duration in our stories (293 ms). There were 3 gaussians in the sequence, occurring at 0 ms, 50 ms and 100 ms relative to feature onset.

Statistical analyses were performed by repeating the simulation procedure 1000 times and shuffling the correspondence between a given “neural generator” (the MEG sensor assigned to the feature) each time that feature was encountered.

All decoding analyses were performed in line with the analysis Methods described above, applied to the empirical MEG data.

## References

1. Gwilliams, L. et al. Computational architecture of speech comprehension in the human brain. Annu. Rev. Linguist. 11, (2024).

2. Mesgarani, N., Cheung, C., Johnson, K. & Chang, E. F. Phonetic feature encoding in human superior temporal gyrus. Science 343, 1006–1010 (2014).

3. Gwilliams, L., King, J.-R., Marantz, A. & Poeppel, D. Neural dynamics of phoneme sequences reveal position-invariant code for content and order. Nat. Commun. 13, 6606 (2022).

4. Oganian, Y. & Chang, E. F. A speech envelope landmark for syllable encoding in human superior temporal gyrus. Sci Adv 5, eaay6279 (2019).

5. Poeppel, D. & Assaneo, M. F. Speech rhythms and their neural foundations. Nat. Rev. Neurosci. 21, 322–334 (2020).

6. Gwilliams, L., Linzen, T., Poeppel, D. & Marantz, A. In Spoken Word Recognition, the Future Predicts the Past. J. Neurosci. 38, 7585–7599 (2018).

7. Gwilliams, L. How the brain composes morphemes into meaning. Philos. Trans. R. Soc. Lond. B Biol. Sci. 375, 20190311 (2020).

8. Keshishian, M. et al. Joint, distributed and hierarchically organized encoding of linguistic features in the human auditory cortex. Nat Hum Behav 7, 740–753 (2023).

9. Bemis, D. K. & Pylkkänen, L. Simple composition: a magnetoencephalography investigation into the comprehension of minimal linguistic phrases. J. Neurosci. 31, 2801–2814 (2011).

10. Pallier, C., Devauchelle, A.-D. & Dehaene, S. Cortical representation of the constituent structure of sentences. Proc. Natl. Acad. Sci. U. S. A. 108, 2522–2527 (2011).

11. Brennan, J. R., Stabler, E. P., Van Wagenen, S. E., Luh, W.-M. & Hale, J. T. Abstract linguistic structure correlates with temporal activity during naturalistic comprehension. Brain Lang. 157-158, 81–94 (2016).

12. Nelson, M. J. et al. Neurophysiological dynamics of phrase-structure building during sentence processing. Proc. Natl. Acad. Sci. U. S. A. 114, E3669–E3678 (2017).

13. Hickok, G. & Poeppel, D. Dorsal and ventral streams: a framework for understanding aspects of the functional anatomy of language. Cognition 92, 67–99 (2004).

14. Hickok, G. & Poeppel, D. The cortical organization of speech processing. Nat. Rev. Neurosci. 8, 393–402 (2007).

15. de Heer, W. A., Huth, A. G., Griffiths, T. L., Gallant, J. L. & Theunissen, F. E. The hierarchical cortical organization of human speech processing. J. Neurosci. 37, 6539–6557 (2017).

16. Federmeier, K. D., Kutas, M. & Dickson, D. A common neural progression to meaning in about a third of a second. Neurobiology of language 557–567 (2015).

17. Newman, R. L. & Connolly, J. F. Electrophysiological markers of pre-lexical speech processing: evidence for bottom-up and top-down effects on spoken word processing. Biol. Psychol. 80, 114–121 (2009).

18. Connolly, J. & Phillips, N. Event-related potential components reflect phonological and semantic processing of the terminal word of spoken sentences. J. Cogn. Neurosci. 6, 256–266 (1994).

19. Desroches, A. S., Newman, R. L. & Joanisse, M. F. Investigating the time course of spoken word recognition: electrophysiological evidence for the influences of phonological similarity. J. Cogn. Neurosci. 21, 1893–1906 (2009).

20. Pylkkänen, L. & Marantz, A. Tracking the time course of word recognition with MEG. Trends Cogn. Sci. 7, 187–189 (2003).

21. Friedrich, C. K., Schild, U. & Röder, B. Electrophysiological indices of word fragment priming allow characterizing neural stages of speech recognition. Biol. Psychol. 80, 105–113 (2009).

22. Friedrich, C. K., Kotz, S. A., Friederici, A. D. & Gunter, T. C. ERPs reflect lexical identification in word fragment priming. J. Cogn. Neurosci. 16, 541–552 (2004).

23. Friedrich, C. K., Kotz, S. A., Friederici, A. D. & Alter, K. Pitch modulates lexical identification in spoken word recognition: ERP and behavioral evidence. Brain Res. Cogn. Brain Res. 20, 300–308 (2004).

24. Kutas, M. & Federmeier, K. D. Thirty years and counting: finding meaning in the N400 component of the event-related brain potential (ERP). Annu. Rev. Psychol. 62, 621–647 (2011).

25. Bornkessel-Schlesewsky, I. & Schlesewsky, M. An alternative perspective on ‘semantic P600’ effects in language comprehension. Brain Res. Rev. 59, 55–73 (2008).

26. Gouvea, A. C., Phillips, C., Kazanina, N. & Poeppel, D. The linguistic processes underlying the P600. Lang. Cogn. Process. 25, 149–188 (2010).

27. Kaan, E., Harris, A., Gibson, E. & Holcomb, P. The P600 as an index of syntactic integration difficulty. Lang. Cogn. Process. 15, 159–201 (2000).

28. Friederici, A., Pfeifer, E. & Hahne, A. Event-related brain potentials during natural speech processing: effects of semantic, morphological and syntactic violations. Brain research. Cognitive brain research 1, 183–192 (1993).

29. Caucheteux, C., Gramfort, A. & King, J.-R. Evidence of a predictive coding hierarchy in the human brain listening to speech. Nat Hum Behav 7, 430–441 (2023).

30. Brennan, J. R. & Hale, J. T. Hierarchical structure guides rapid linguistic predictions during naturalistic listening. PLoS One 14, e0207741 (2019).

31. Brodbeck, C., Hong, L. E. & Simon, J. Z. Rapid Transformation from Auditory to Linguistic Representations of Continuous Speech. Curr. Biol. 28, 3976–3983.e5 (2018).

32. D’efossez, A., Caucheteux, C., Rapin, J., Kabeli, O. & King, J. Decoding speech perception from non-invasive brain recordings. Nat. Mach. Intell. 5, 1097–1107 (2022).

33. Heilbron, M., Armeni, K., Schoffelen, J.-M., Hagoort, P. & de Lange, F. P. A hierarchy of linguistic predictions during natural language comprehension. Proc. Natl. Acad. Sci. U. S. A. 119, e2201968119 (2022).

34. Honey, C. J. et al. Slow cortical dynamics and the accumulation of information over long timescales. Neuron 76, 423–434 (2012).

35. Chen, C., Dupré la Tour, T., Gallant, J. L., Klein, D. & Deniz, F. The cortical representation of language timescales is shared between reading and listening. Commun Biol 7, 284 (2024).

36. Lerner, Y., Honey, C. J., Silbert, L. J. & Hasson, U. Topographic mapping of a hierarchy of temporal receptive windows using a narrated story. J. Neurosci. 31, 2906–2915 (2011).

37. Connine, C., Blasko, D. G. & Hall, M. D. Effects of subsequent sentence context in auditory word recognition: Temporal and linguistic constrainst. Journal of Memory and Language 30, 234–250 (1991).

38. Bicknell, K., Bushong, W., Tanenhaus, M. K. & Jaeger, T. F. Maintenance of subcategorical information during speech perception: revisiting misunderstood limitations. J. Mem. Lang. 140, 104565 (2025).

39. Levy, R., Bicknell, K., Slattery, T. & Rayner, K. Eye movement evidence that readers maintain and act on uncertainty about past linguistic input. Proc. Natl. Acad. Sci. U. S. A. 106, 21086–21090 (2009).

40. Levinson, S. ‘Process and perish’ or multiple buffers with push-down stacks? [Commentary on The Now-or-Never Bottleneck: A Fundamental Constraint on Language by M.H. Christiansen and N. Chater]. Behavioral and Brain Sciences (2015).

41. Fedorenko, E. & Thompson-Schill, S. L. Reworking the language network. Trends Cogn. Sci. 18, 120–126 (2014).

42. Desbordes, T. et al. Dimensionality and Ramping: Signatures of Sentence Integration in the Dynamics of Brains and Deep Language Models. J. Neurosci. 43, 5350–5364 (2023).

43. King, J.-R. & Dehaene, S. Characterizing the dynamics of mental representations: the temporal generalization method. Trends Cogn. Sci. 18, 203–210 (2014).

44. Stokes, M. G., Buschman, T. J. & Miller, E. K. Dynamic coding for flexible cognitive control. in The Wiley Handbook of Cognitive Control 221–241 (John Wiley & Sons, Ltd, Chichester, UK, 2017).

45. Stroud, J. P., Watanabe, K., Suzuki, T., Stokes, M. G. & Lengyel, M. Optimal information loading into working memory explains dynamic coding in the prefrontal cortex. Proc. Natl. Acad. Sci. U. S. A. 120, e2307991120 (2023).

46. King, J.-R. & Wyart, V. The Human Brain Encodes a Chronicle of Visual Events at Each Instant of Time Through the Multiplexing of Traveling Waves. J. Neurosci. 41, 7224–7233 (2021).

47. Brennan, J. & Pylkkänen, L. The time-course and spatial distribution of brain activity associated with sentence processing. Neuroimage 60, 1139–1148 (2012).

48. Bicknell, K., Jaeger, T. F. & Tanenhaus, M. K. Now or … later: Perceptual data are not immediately forgotten during language processing. Behav. Brain Sci. 39, e67 (2016).

49. van Berkum, J. J., Hagoort, P. & Brown, C. M. Semantic integration in sentences and discourse: evidence from the N400. J. Cogn. Neurosci. 11, 657–671 (1999).

50. Huth, A. G., de Heer, W. A., Griffiths, T. L., Theunissen, F. E. & Gallant, J. L. Natural speech reveals the semantic maps that tile human cerebral cortex. Nature 532, 453–458 (2016).

51. Su, J. et al. RoFormer: Enhanced transformer with Rotary Position Embedding. Neurocomputing 568, 127063 (2024).

52. Peng, B., Quesnelle, J., Fan, H. & Shippole, E. YaRN: Efficient Context Window Extension of Large Language Models. arXiv [cs.CL] (2023).

53. Ding, N. Sequence chunking through neural encoding of ordinal positions. Trends Cogn. Sci. (2025) doi:10.1016/j.tics.2025.01.014.

54. Caucheteux, C., Gramfort, A. & King, J.-R. Model-based analysis of brain activity reveals the hierarchy of language in 305 subjects. arXiv [q-bio.NC] (2021).

55. Jain, S., Vo, V. A., Wehbe, L. & Huth, A. G. Computational language modeling and the promise of in silico experimentation. Neurobiology of Language 1–65 (2023).

56. Chang, C. H. C., Nastase, S. A. & Hasson, U. Information flow across the cortical timescale hierarchy during narrative construction. Proc. Natl. Acad. Sci. U. S. A. 119, e2209307119 (2022).

57. Levy, R. Communicative efficiency, Uniform Information Density, and the Rational Speech Act theory. CogSci (2018) doi:10.31234/osf.io/4cgxh.

58. Gwilliams, L. & King, J.-R. Recurrent processes support a cascade of hierarchical decisions. Elife 9, (2020).

59. Abrams, E. B., Marantz, A., Krementsov, I. & Gwilliams, L. Dynamics of pitch perception in the auditory cortex. J. Neurosci. e1111242025 (2025).

60. Lau, E. F., Phillips, C. & Poeppel, D. A cortical network for semantics: (de)constructing the N400. Nat. Rev. Neurosci. 9, 920–933 (2008).

61. Kutas, M. & Federmeier, K. D. Electrophysiology reveals semantic memory use in language comprehension. Trends Cogn. Sci. 4, 463–470 (2000).

62. Hagoort, P., Hald, L., Bastiaansen, M. & Petersson, K. M. Integration of word meaning and world knowledge in language comprehension. Science 304, 438–441 (2004).

63. Lau, E. F., Namyst, A. M., Fogel, A. & Delgado, T. A direct comparison of N400 effects of predictability and incongruity in adjective-noun combination. Collabra 2, (2016).

64. Kuperberg, G. R. & Jaeger, T. F. What do we mean by prediction in language comprehension? Lang. Cogn. Neurosci. 31, 32–59 (2016).

65. Wang, L., Brothers, T., Jensen, O. & Kuperberg, G. R. Dissociating the pre-activation of word meaning and form during sentence comprehension: Evidence from EEG representational similarity analysis. Psychon. Bull. Rev. 31, 862–873 (2024).

66. Hubbard, R. J. & Federmeier, K. D. The impact of linguistic prediction violations on downstream recognition memory and sentence recall. J. Cogn. Neurosci. 1–23 (2023).

67. Gwilliams, L., Marantz, A., Poeppel, D. & King, J.-R. Top-down information shapes lexical processing when listening to continuous speech. Lang. Cogn. Neurosci. 39, 1045–1058 (2024).

68. Ferreira, F., Bailey, K. G. D. & Ferraro, V. Good-Enough Representations in Language Comprehension. Curr. Dir. Psychol. Sci. 11, 11–15 (2002).

69. Frances, C. Good enough processing: what have we learned in the 20 years since Ferreira et al. (2002)? Front. Psychol. 15, (2024).

70. Ferreira, F. & Patson, N. D. The ‘good enough’ approach to language comprehension: The ‘good enough’ approach. Lang. Linguist. Compass 1, 71–83 (2007).

71. Friederici, A. D. Towards a neural basis of auditory sentence processing. Trends Cogn. Sci. 6, 78–84 (2002).

72. Hochstein, S. & Ahissar, M. View from the top: hierarchies and reverse hierarchies in the visual system. Neuron 36, 791–804 (2002).

73. Ahissar, M. & Hochstein, S. The reverse hierarchy theory of visual perceptual learning. Trends Cogn. Sci. 8, 457–464 (2004).

74. Warren, R. M. Perceptual restoration of missing speech sounds. Science 167, 392–393 (1970).

75. Leonard, M. K., Baud, M. O., Sjerps, M. J. & Chang, E. F. Perceptual restoration of masked speech in human cortex. Nat. Commun. 7, 13619 (2016).

76. Luke, S. G. & Christianson, K. Limits on lexical prediction during reading. Cogn. Psychol. 88, 22–60 (2016).

77. Davis, M. H. & Johnsrude, I. S. Hearing speech sounds: top-down influences on the interface between audition and speech perception. Hear. Res. 229, 132–147 (2007).

78. Cope, T. E. et al. Evidence for causal top-down frontal contributions to predictive processes in speech perception. Nat. Commun. 8, 2154 (2017).

79. Nahum, M., Nelken, I. & Ahissar, M. Low-level information and high-level perception: the case of speech in noise. PLoS Biol. 6, e126 (2008).

80. Gwilliams, L. et al. Introducing MEG-MASC a high-quality magneto-encephalography dataset for evaluating natural speech processing. Sci Data 10, 862 (2023).

81. Ide, N. & Macleod, C. The american national corpus: A standardized resource of american english. in Proceedings of corpus linguistics vol. 3 1–7 (Lancaster University Centre for Computer Corpus Research on Language …, 2001).

82. Yuan, J. & Liberman, M. Speaker identification on the SCOTUS corpus. J. Acoust. Soc. Am. 123, 3878–3878 (2008).

83. Fedorenko, E., Ivanova, A. A. & Regev, T. I. The language network as a natural kind within the broader landscape of the human brain. Nat. Rev. Neurosci. (2024) doi:10.1038/s41583-024-00802-4.

84. Ochshorn, R. M. & Hawkins, M. Gentle forced aligner. github. com/lowerquality/gentle (2017).

85. Adachi, Y., Shimogawara, M., Higuchi, M., Haruta, Y. & Ochiai, M. Reduction of non-periodic environmental magnetic noise in MEG measurement by continuously adjusted least squares method. IEEE Trans. Appl. Supercond. 11, 669–672 (2001).

86. Gramfort, A. et al. MNE software for processing MEG and EEG data. Neuroimage 86, 446–460 (2014).

87. King, S. & Taylor, P. Detection of phonological features in continuous speech using neural networks. Comput. Speech Lang. 14, 333–353 (2000).

88. Balota, D. A. et al. The English Lexicon Project. Behav. Res. Methods 39, 445–459 (2007).

89. King, J.-R., Charton, F., Lopez-Paz, D. & Oquab, M. Back-to-back regression: Disentangling the influence of correlated factors from multivariate observations. Neuroimage 220, 117028 (2020).

90. King, J.-R., Gramfort, A. & Others. Encoding and decoding neuronal dynamics: Methodological framework to uncover the algorithms of cognition. (2018).

91. Pedregosa, F. et al. Scikit-learn: Machine Learning in Python. arXiv [cs.LG] 2825–2830 (2012).

