## Supplement for "Hierarchical dynamic coding coordinates speech comprehension in the human brain"

### 1. Supplementary Results

#### *1.1. Distribution of word sequences across time*

We observed that language features are maintained for a long time, potentially overlapping with the processing of many words into the future. To quantify the extent of this overlap, we computed the timing distribution of words in our stories.

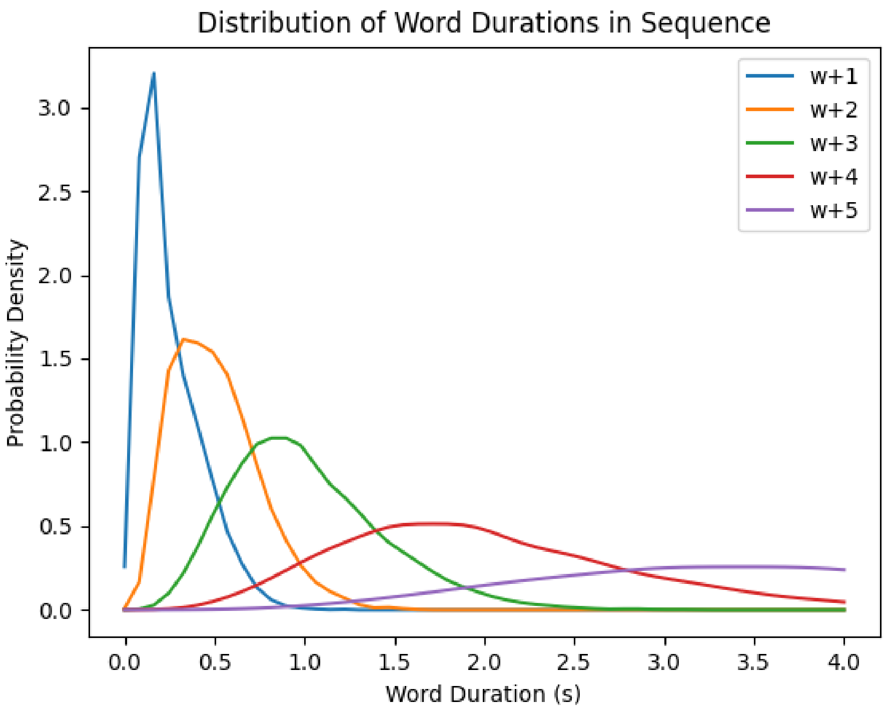


Figure 1: **Distribution of word duration.** Probability density function for the word onsets of the words 1, 2, 3, 4, and 5 words into the future sequence. This demonstrates that when a feature of w0 is maintained for 1 second into the future, this is, on average, being maintained into the processing of three or four words into the future.

*1.2. Hierarchy decoding at word onset*

We repeated the same analysis displayed in Main Figure 4A time-locked to word onset. The results of the permutation test are quantitatively weaker, but qualitatively comparable to the offset results. Phonetic features can be decoded from around 112:368 ms ($\hat{t}$ (average $t$-value in the cluster) = 1.96, $p$ = .048); sub-lexical features from 80:304 ms ($\hat{t}$ = 1.56, $p$ = .15); word class from 30:424 ms ($\hat{t}$ = 2.14, $p$ = .013); syntactic operation from 232:944 ms ($\hat{t}$ = 2.33, $p$ = .004); syntactic state ($\hat{t}$ = 3.57, $p$ < .001) and semantic vectors ($\hat{t}$ = 3.97, $p$ < .001) throughout the entire search window.

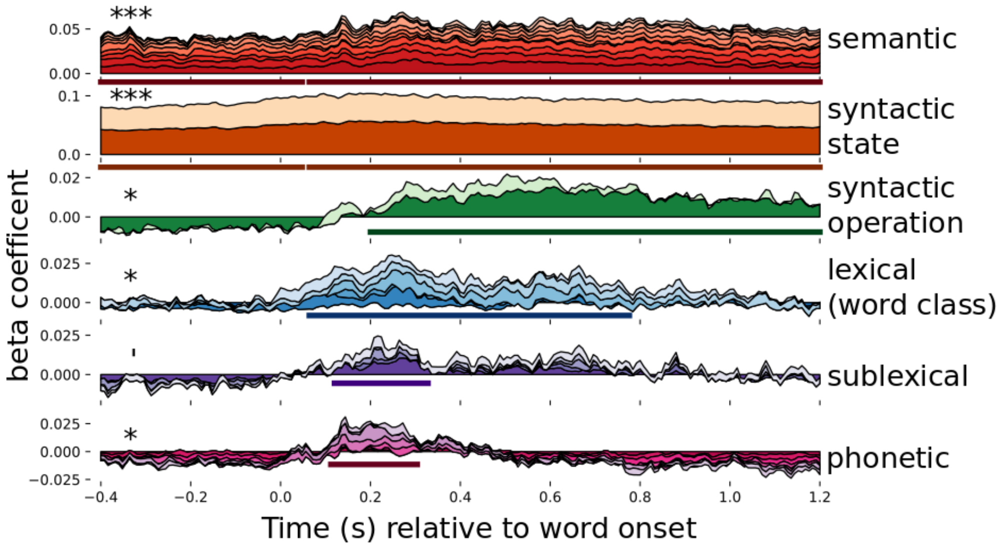


Figure 2: **Word onset language level decoding.** Result of decoding each language level over time when time locking the analysis to word onset. The beta coefficients of each feature are stacked on top of each other, such that the top of the timecourse plot corresponds to the cumulative sum of all features in that linguistic level. The x-axis corresponds to time in seconds relative to word offset. The y-axis corresponds to the cumulative beta-coefficient across features. Solid line below the time-course represents the extent of the significant temporal cluster; asterisks represent its significance: * p < .05; ** p < .01; *** p < .001.

#### *1.3. Temporal Generalization Analysis Locked to Word-Onset*

We repeated the same analysis displayed in Main Figure 5A time-locked to word onset. Decoding performance is numerically weaker, but the generalization pattern is similar to the word offset results: Wider diagonal pattern for increasing levels of the language hiearchy.

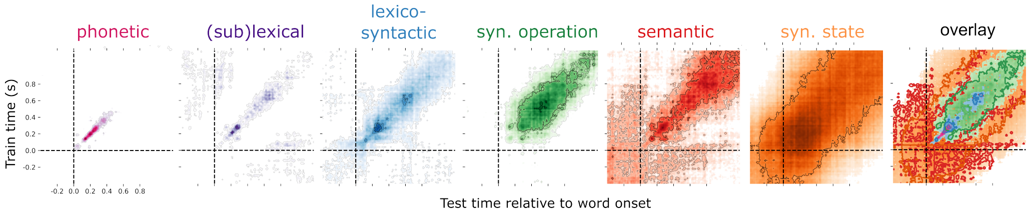


Figure 3: **Word onset temporal generalization.** Result of temporal generalization decoding at each language level over time when time locking the analysis to word onset.

*1.4. Replication across sessions*

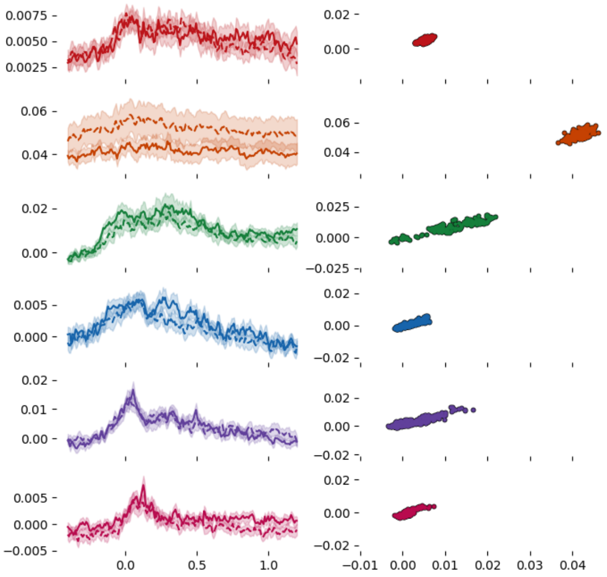


Figure 4: **Replicability across sessions.** Left: Average decoding accuracy is displayed for each of the six linguistic levels, from session one (dashed line) and session two (solid line). Right: Correlation between the average timecourses across subjects.

#### 1.5. *Detailed decoding results for each linguistic feature*

If the decoding timecourses of a particular feature are significantly different from one another, we can use this as evidence for separable underlying processes. The results of this decoding time-locked to word offset is presented in Supplementary Figure 5, and relative to word onset is presented in Supplementary Figure 6 and displayed in each of the tables below.

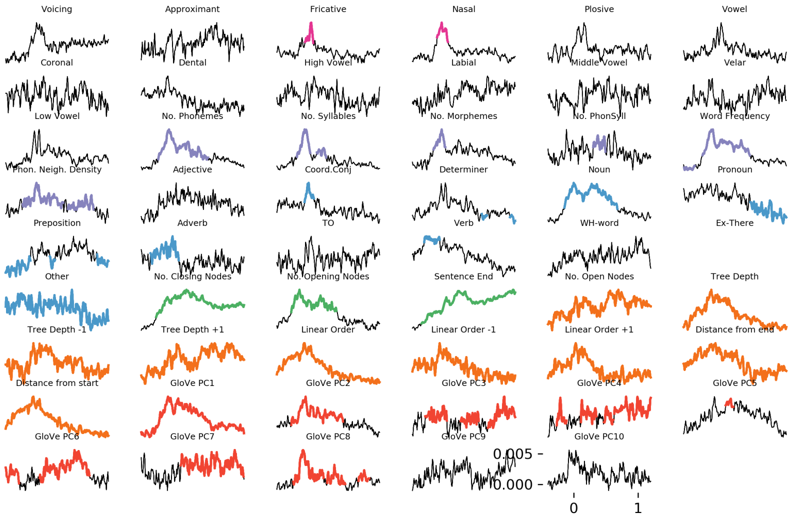


Figure 5: **Word offset feature decoding.** Average R-values across subjects for each of the linguistic features of interest, time-locked to word offset. Coloured portions of the time course correspond to temporal clusters that exceed a $p < .05$ threshold. Colour corresponds to the putative linguistic level.

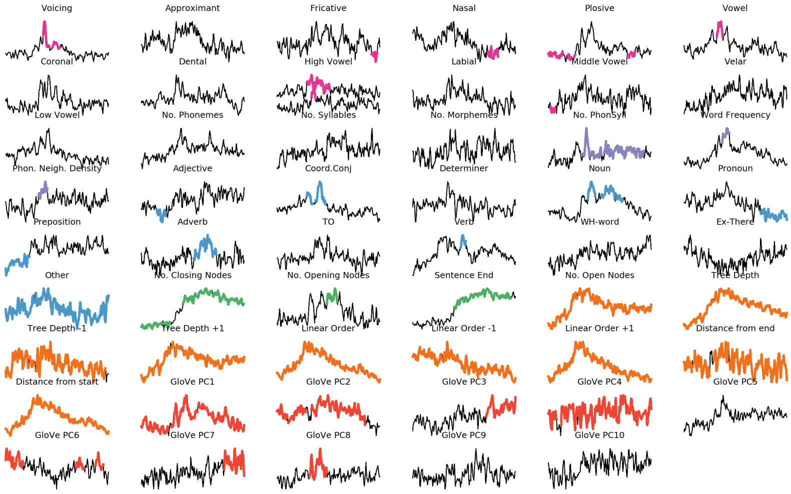


Figure 6: **Word onset feature decoding.** Average R-values across subjects for each of the linguistic features of interest, time-locked to word onset. Coloured portions of the time course correspond to temporal clusters that exceed a $p < .05$ threshold. Colour corresponds to the putative linguistic level.

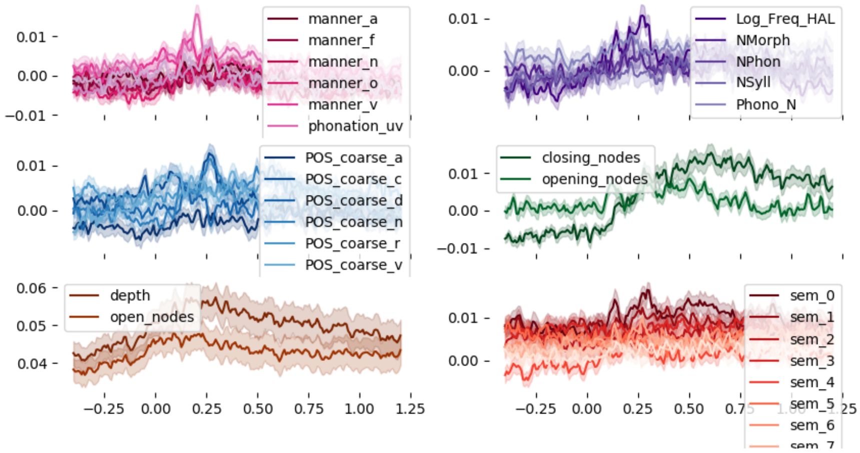


Figure 7: **Word onset feature decoding grouped.** Average R-values across subjects for each of the linguistic features of interest, time-locked to word onset, and grouped into feature families as shown in Figure 3. Coloured portions of the time course correspond to temporal clusters that exceed a $p < .05$ threshold. Colour corresponds to the putative linguistic level.

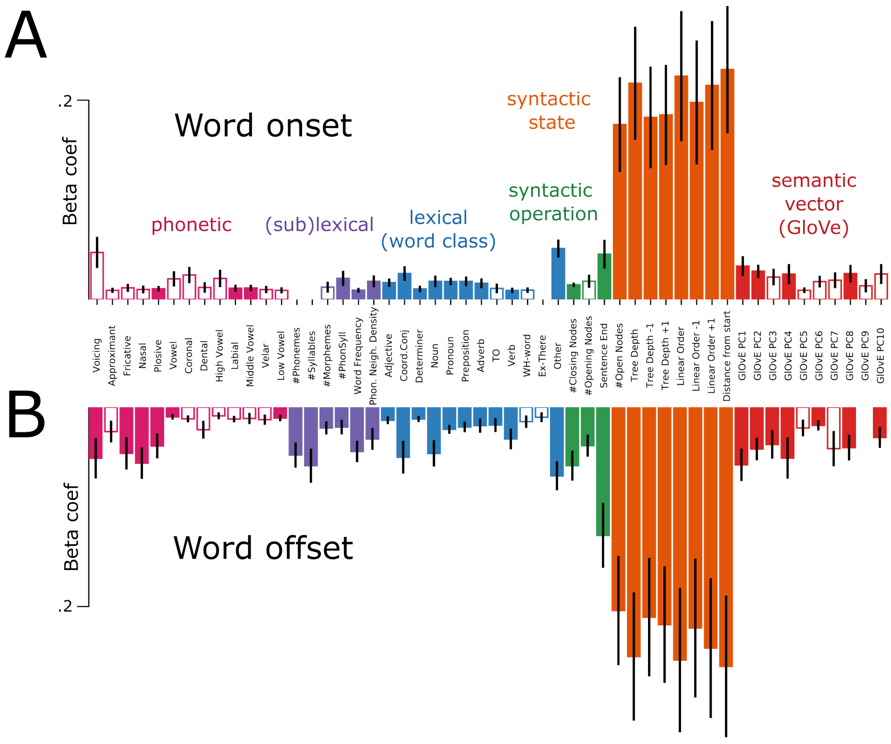


Figure 8: **Word onset and word offset average decoding performance.** Barplots represent average decoding over time, and error bars represent standard error of the mean. Filled bars correspond to features that are decodable significantly better than chance at p ¡ .05 level, confirmed with a random permutation test.

#### *1.6. Simulations*

In order to test the assumptions of the HDC hypothesis, we simulated the MEG data under different conditions. Below are the results of our simulations replicating main Figure 4.

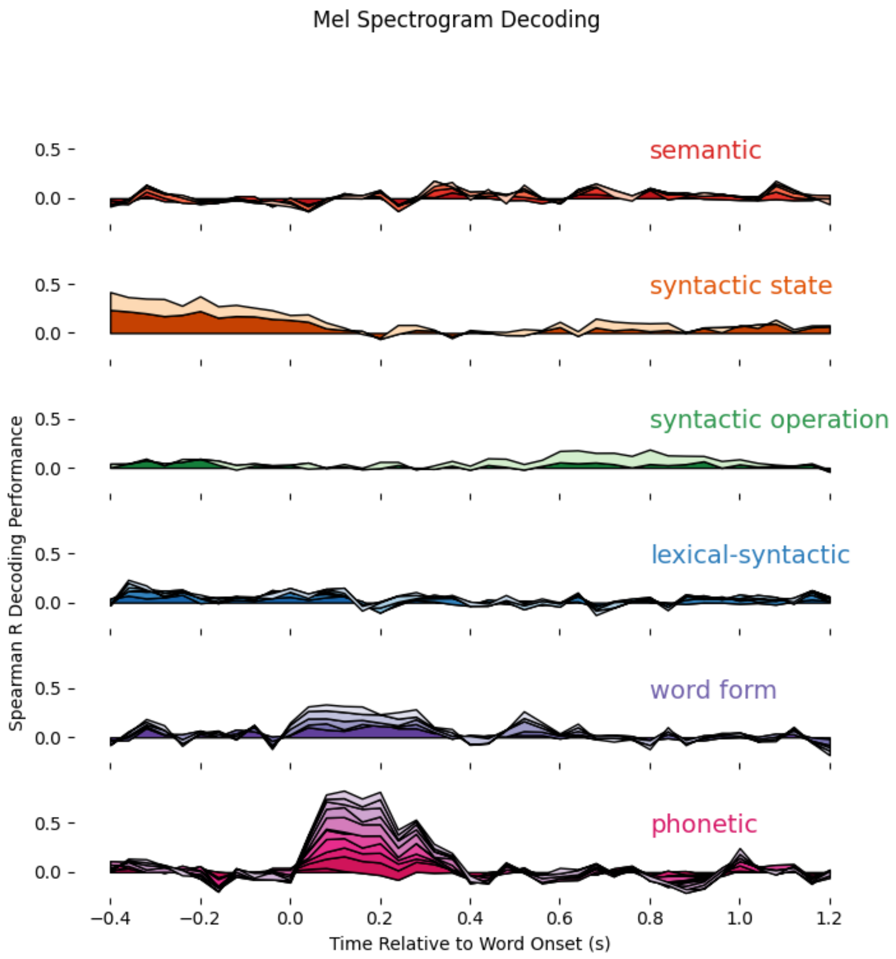


Figure 9: **Decoding language hierarchy from mel spectrogram.** The beta coefficients of each feature are stacked on top of each other, such that the top of the timecourse plot corresponds to the cumulative sum of all features in that linguistic level. The x-axis corresponds to time in seconds relative to word offset. The y-axis corresponds to the cumulative beta-coefficient across features.

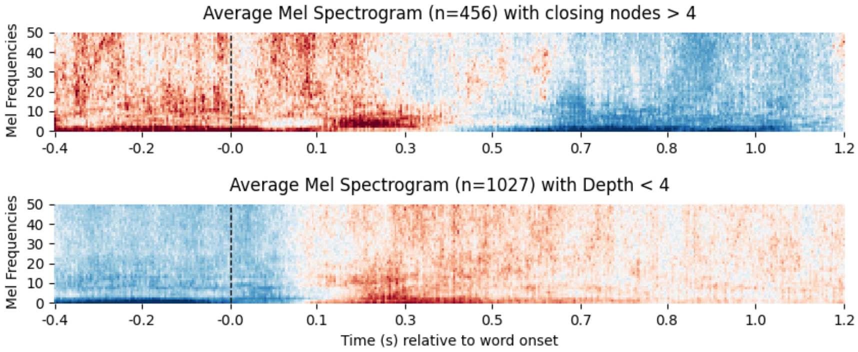


Figure 10: **Average Mel spectrogram for different language features.** We observed above-chance decoding for syntactic features from the Mel spectrogram, so we wanted to investigate the acoustic origin. We found that a large number of closing nodes are likely to occur at a sentence offset, thus leading to acoustic transitions into silence (above). Similarly, shallow sentence depth often occurs towards the beginning of a sentence, leading to systematic acoustic onsets (below).

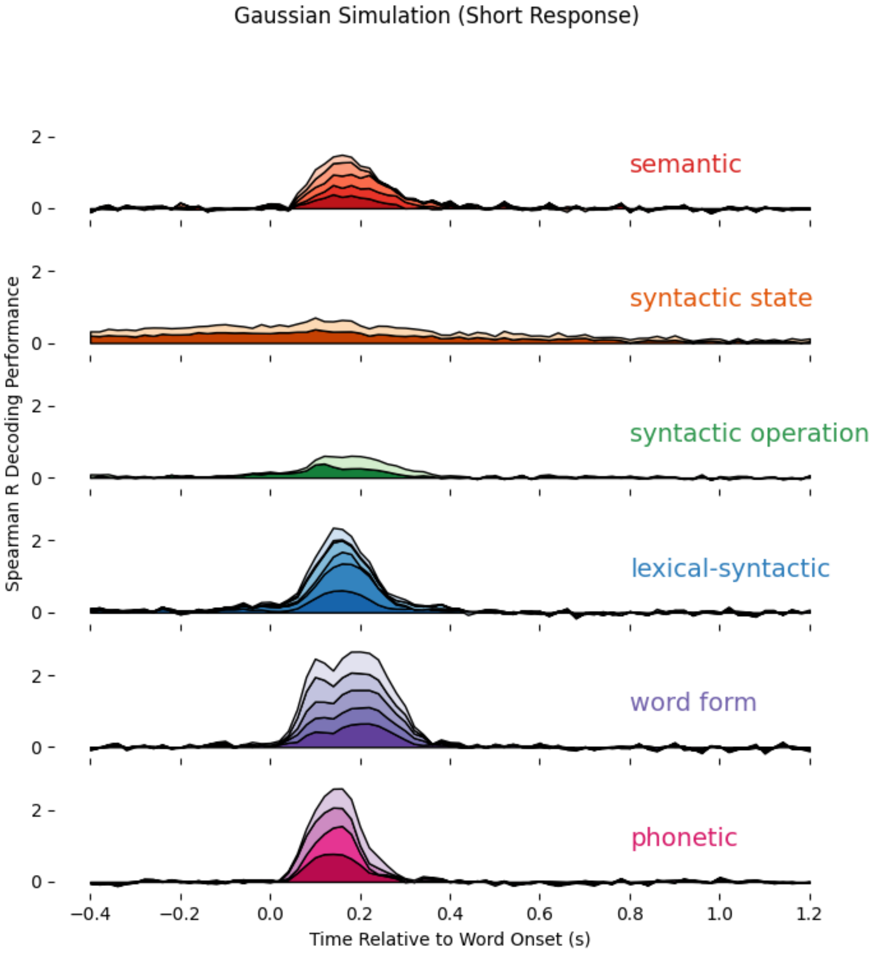


Figure 11: **Simulating MEG data as gaussian responses to language input.** The beta coefficients of each feature are stacked on top of each other, such that the top of the timecourse plot corresponds to the cumulative sum of all features in that linguistic level. The x-axis corresponds to time in seconds relative to word offset. The y-axis corresponds to the cumulative beta-coefficient across features.

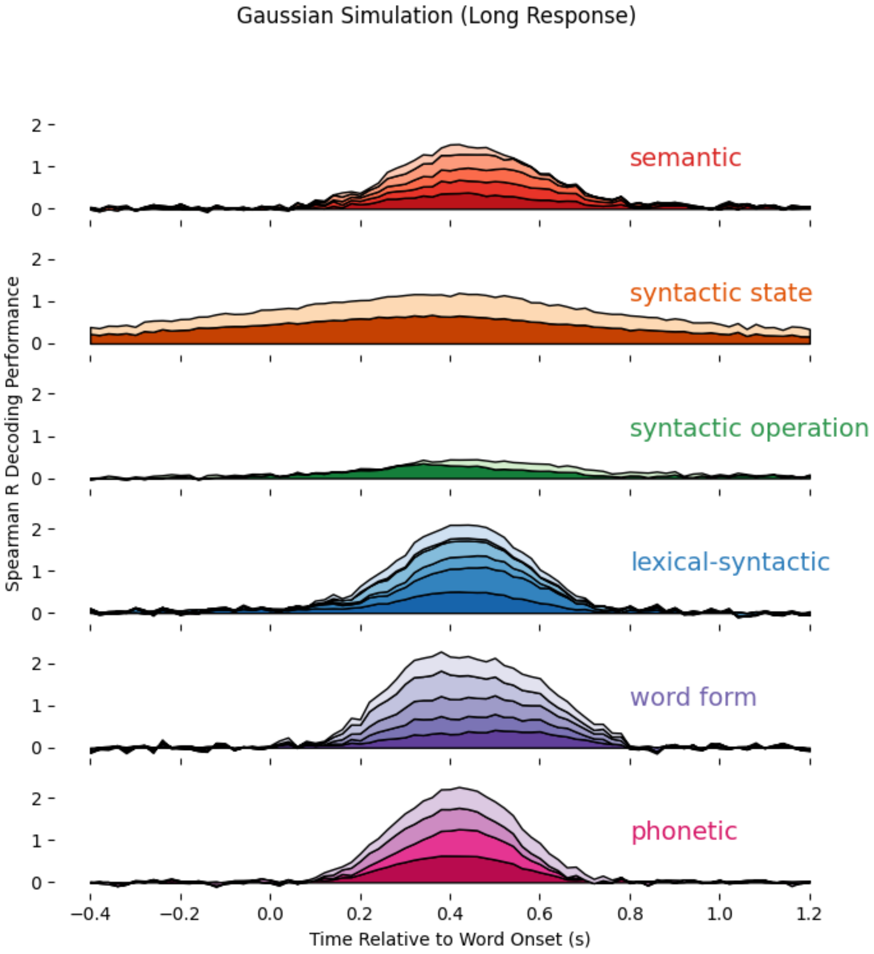


Figure 12: **Simulating MEG data as prolonged gaussian responses to language input.** The beta coefficients of each feature are stacked on top of each other, such that the top of the timecourse plot corresponds to the cumulative sum of all features in that linguistic level. The x-axis corresponds to time in seconds relative to word offset. The y-axis corresponds to the cumulative beta-coefficient across features.

*1.7. Evoked feature responses*

To connect our results to the comprehensive body of literature using ERP-style analyses, we also used regression encoding models to reconstruct the sensor-level responses to each of our stimulus features. Below we provide the full sensor response time-locked to word offset, as well as the root-mean-square of responses, when conducting median split of the trials.

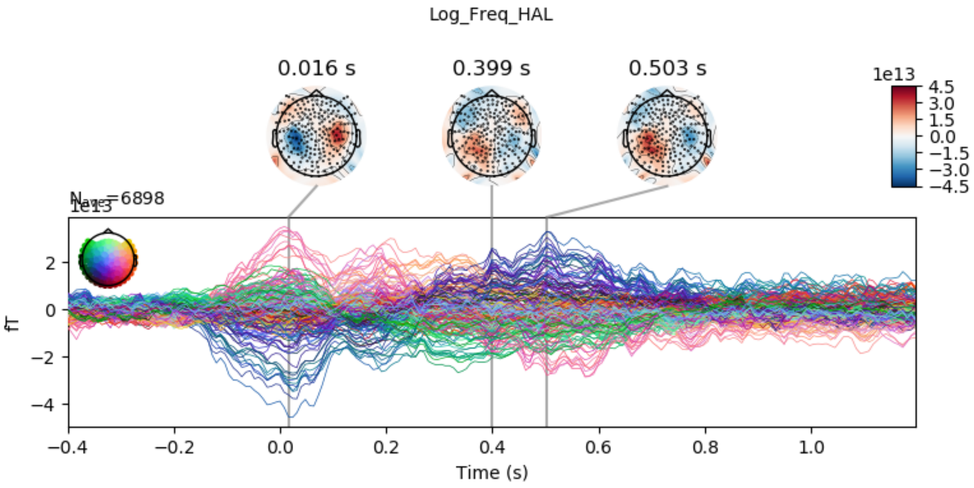


Figure 13: **Log Word Frequency sensor encoding.** x-axis represents time in seconds relative to word offset; y-axis represents the femto-tesla activity strength of each sensor.

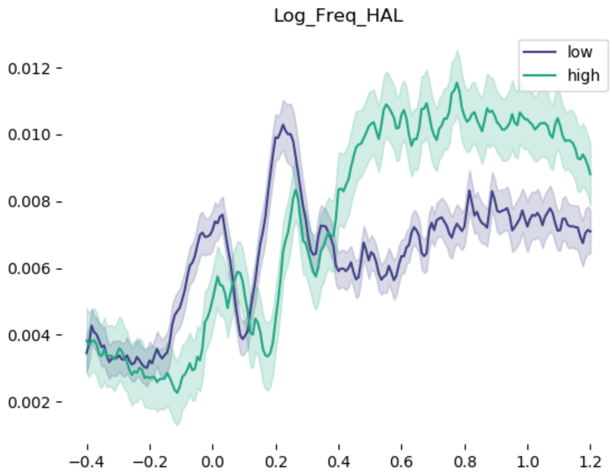


Figure 14: **Log Word Frequency RMS median split.** x-axis represents time in seconds relative to word offset; y-axis represents the root-mean-square of femto-tesla activity strength over all sensors.

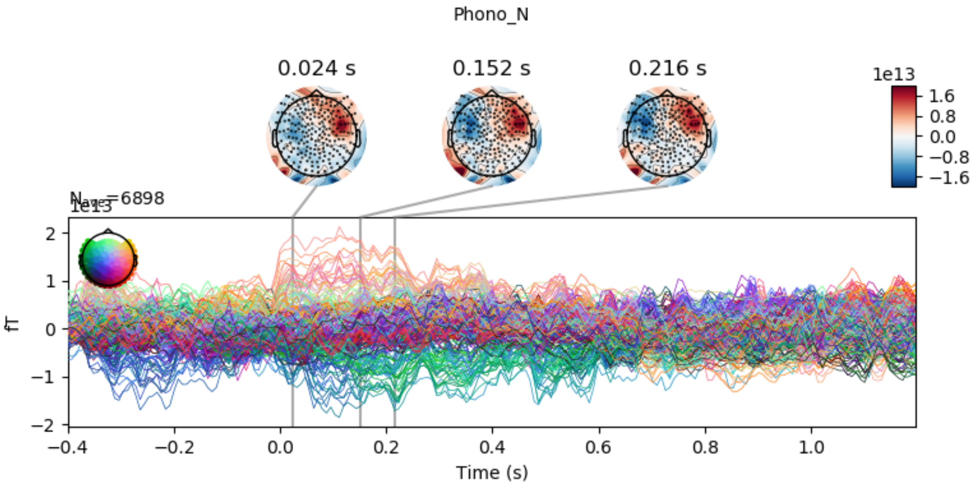


Figure 15: **Phonological Neighbourhood Density sensor encoding.** x-axis represents time in seconds relative to word offset; y-axis represents the femto-tesla activity strength of each sensor.

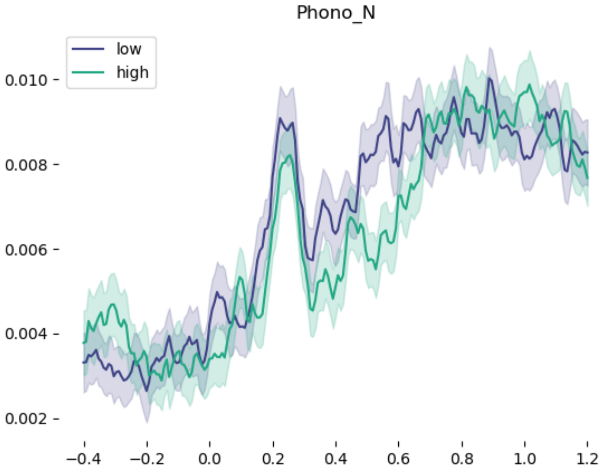


Figure 16: **Phonological Neighbourhood Density RMS median split.** x-axis represents time in seconds relative to word offset; y-axis represents the root-mean-square of femto-tesla activity strength over all sensors.

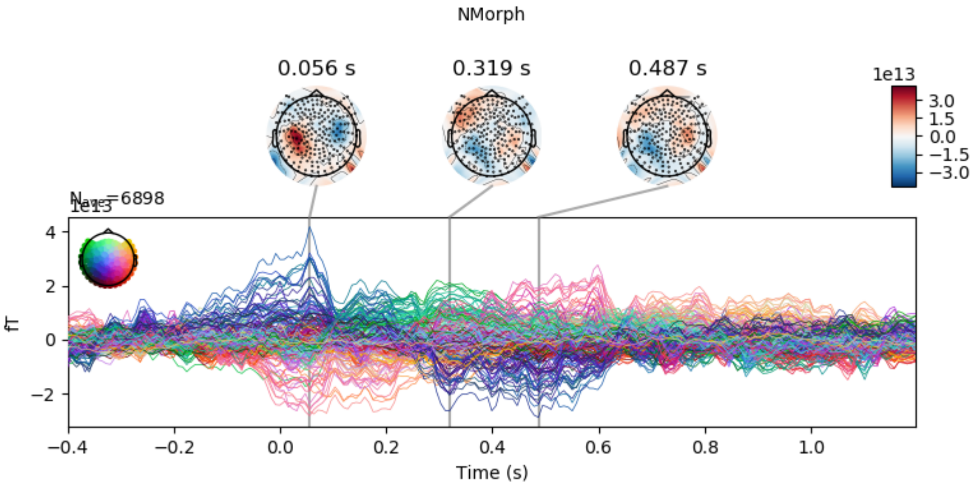


Figure 17: **Number of Morphemes sensor encoding.** x-axis represents time in seconds relative to word offset; y-axis represents the femto-tesla activity strength of each sensor.

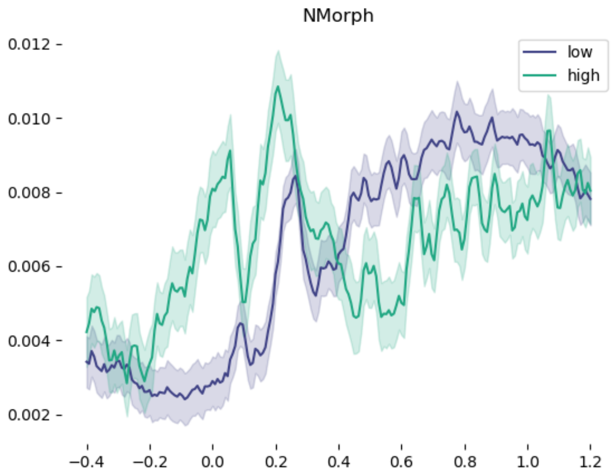


Figure 18: **Number of Morphemes RMS median split.** x-axis represents time in seconds relative to word offset; y-axis represents the root-mean-square of femto-tesla activity strength over all sensors.

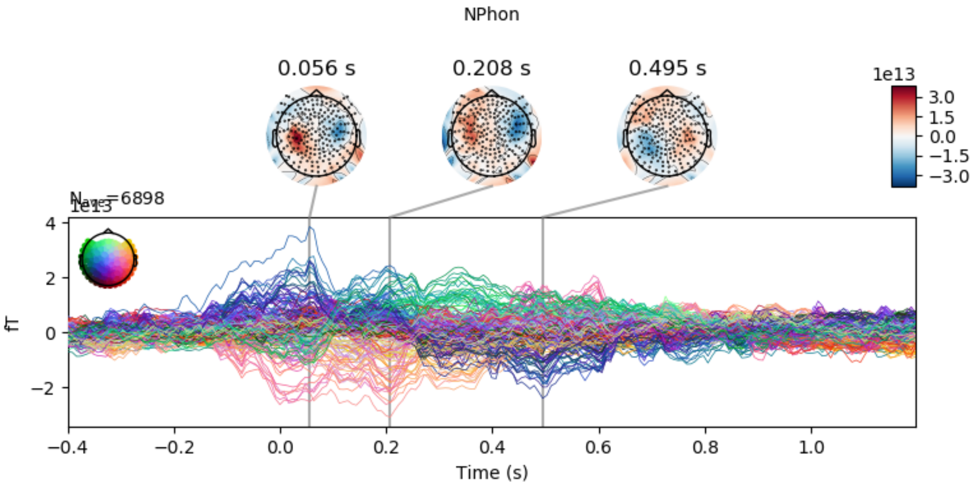


Figure 19: **Number of Phonemes sensor encoding.** x-axis represents time in seconds relative to word offset; y-axis represents the femto-tesla activity strength of each sensor.

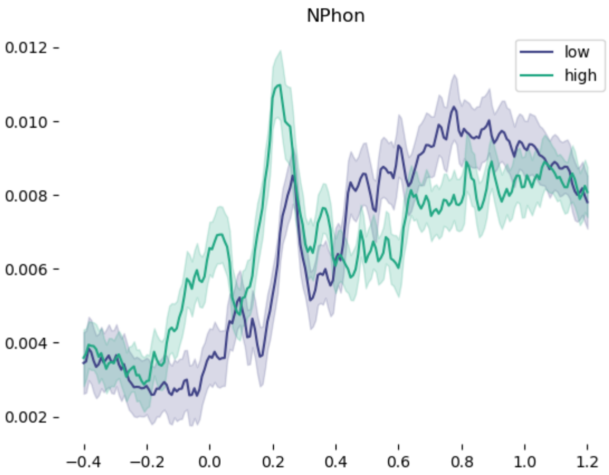


Figure 20: **Number of Phonemes RMS median split.** x-axis represents time in seconds relative to word offset; y-axis represents the root-mean-square of femto-tesla activity strength over all sensors.

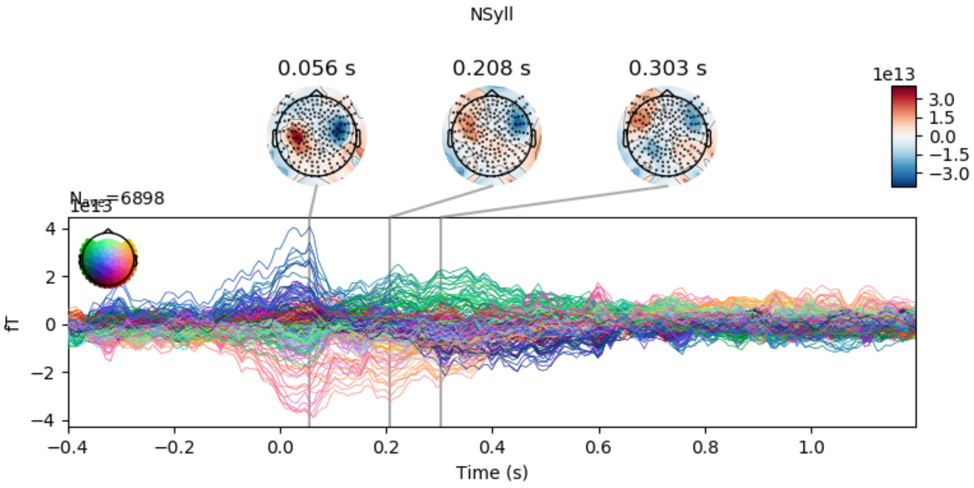


Figure 21: **Number of Syllables sensor encoding.** x-axis represents time in seconds relative to word offset; y-axis represents the femto-tesla activity strength of each sensor.

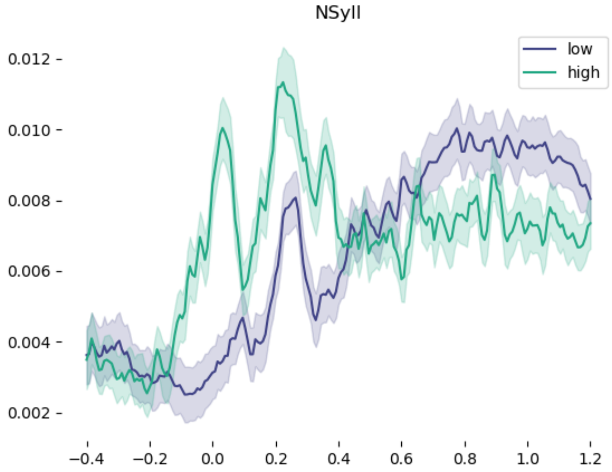


Figure 22: **Number of Syllables RMS median split.** x-axis represents time in seconds relative to word offset; y-axis represents the root-mean-square of femto-tesla activity strength over all sensors.

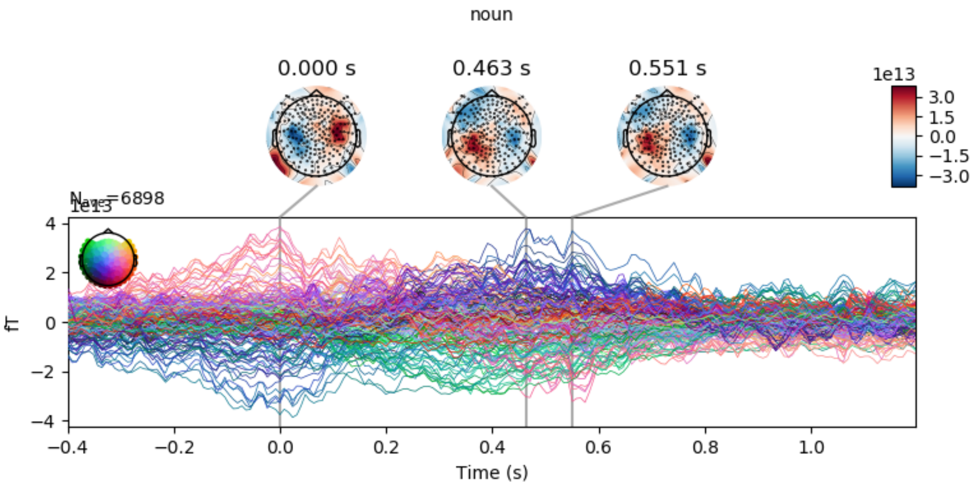


Figure 23: **Noun part-of-speech sensor encoding.** x-axis represents time in seconds relative to word offset; y-axis represents the femto-tesla activity strength of each sensor.

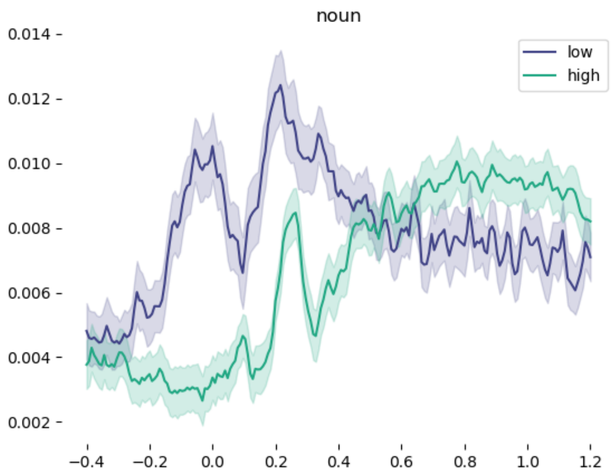


Figure 24: **Noun part-of-speech RMS median split.** x-axis represents time in seconds relative to word offset; y-axis represents the root-mean-square of femto-tesla activity strength over all sensors.

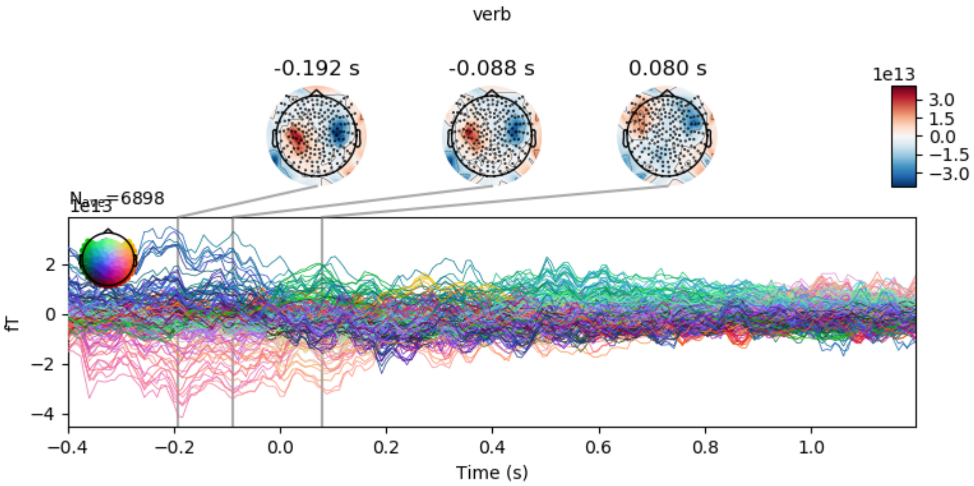


Figure 25: **Verb part-of-speech sensor encoding.** x-axis represents time in seconds relative to word offset; y-axis represents the femto-tesla activity strength of each sensor.

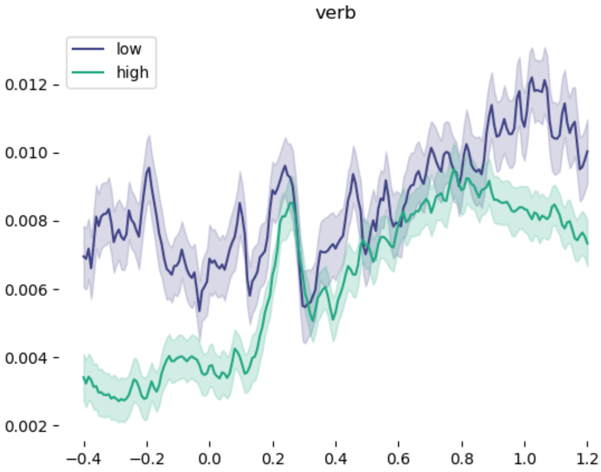


Figure 26: **Verb part-of-speech RMS median split.** x-axis represents time in seconds relative to word offset; y-axis represents the root-mean-square of femto-tesla activity strength over all sensors.

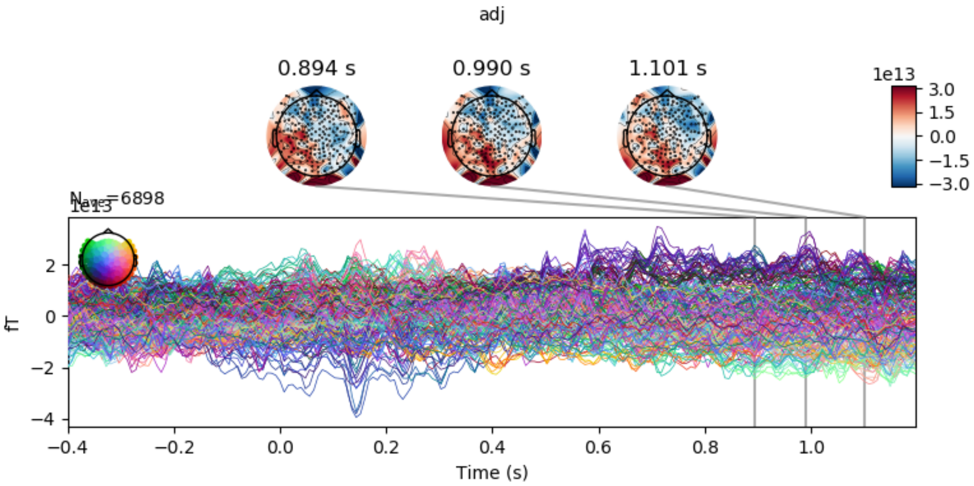


Figure 27: **Adjective part-of-speech sensor encoding.** x-axis represents time in seconds relative to word offset; y-axis represents the femto-tesla activity strength of each sensor.

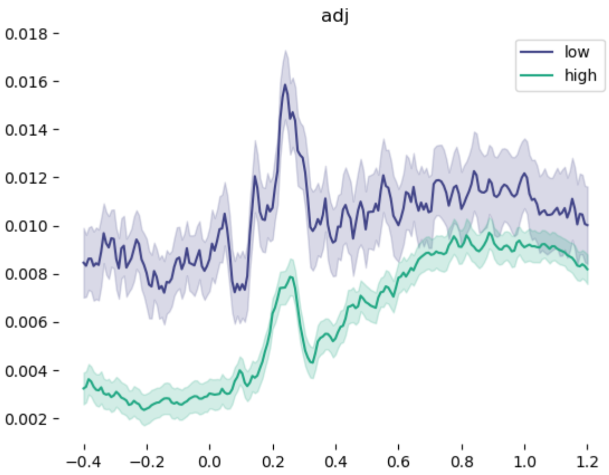


Figure 28: **Adjective part-of-speech RMS median split.** x-axis represents time in seconds relative to word offset; y-axis represents the root-mean-square of femto-tesla activity strength over all sensors.

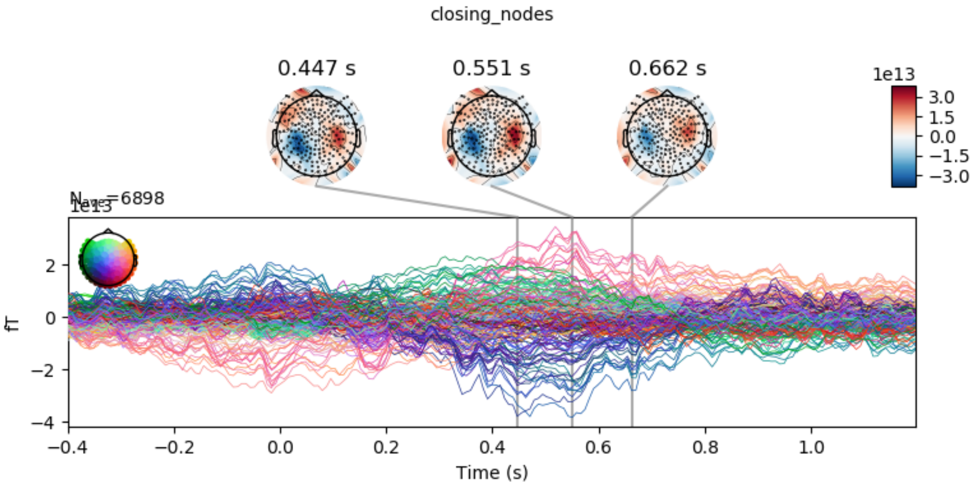


Figure 29: **Number of closing nodes sensor encoding.** x-axis represents time in seconds relative to word offset; y-axis represents the femto-tesla activity strength of each sensor.

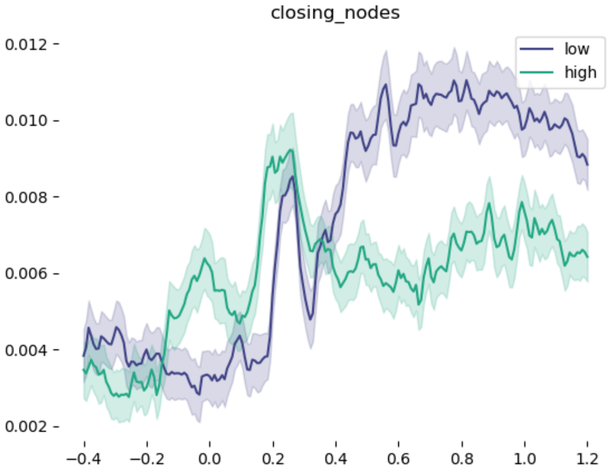


Figure 30: **Number of closing nodes RMS median split.** x-axis represents time in seconds relative to word offset; y-axis represents the root-mean-square of femto-tesla activity strength over all sensors.

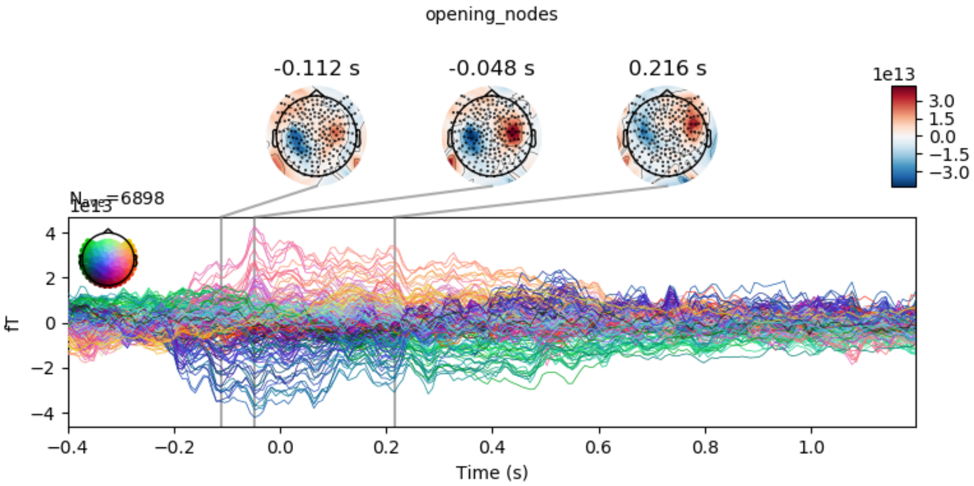


Figure 31: **Number of opening nodes sensor encoding.** x-axis represents time in seconds relative to word offset; y-axis represents the femto-tesla activity strength of each sensor.

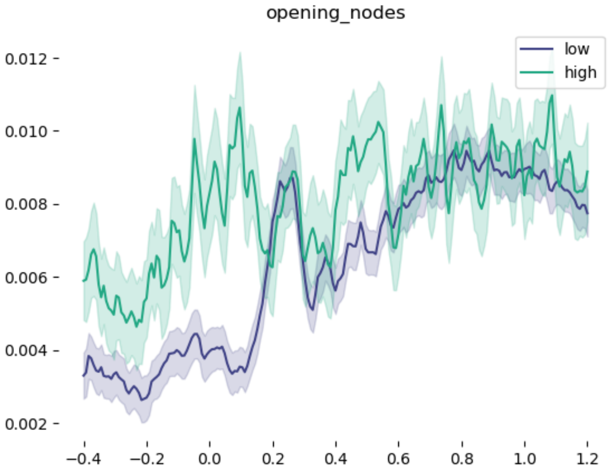


Figure 32: **Number of opening nodes RMS median split.** x-axis represents time in seconds relative to word offset; y-axis represents the root-mean-square of femto-tesla activity strength over all sensors.

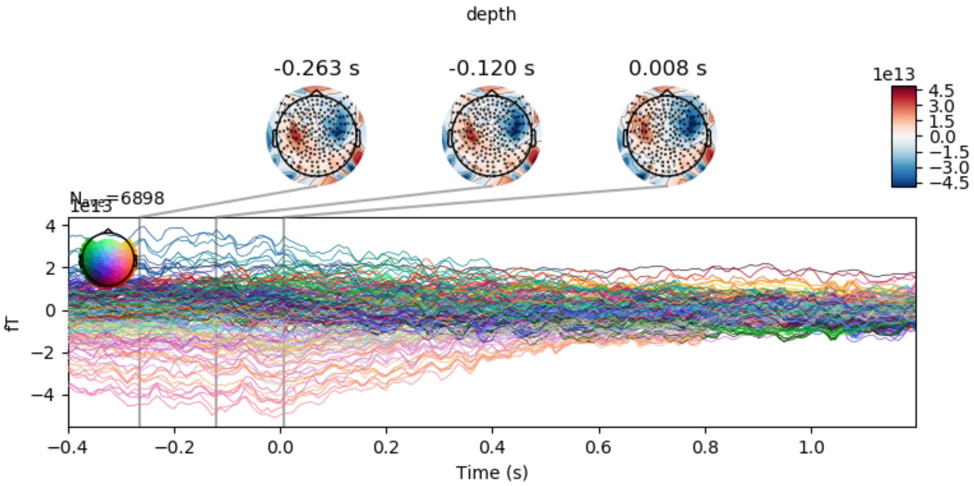


Figure 33: **Tree depth sensor encoding.** x-axis represents time in seconds relative to word offset; y-axis represents the femto-tesla activity strength of each sensor.

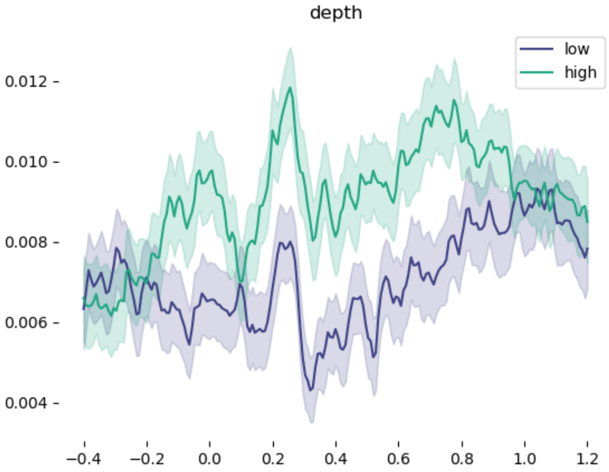


Figure 34: **Tree depth RMS median split.** x-axis represents time in seconds relative to word offset; y-axis represents the root-mean-square of femto-tesla activity strength over all sensors.

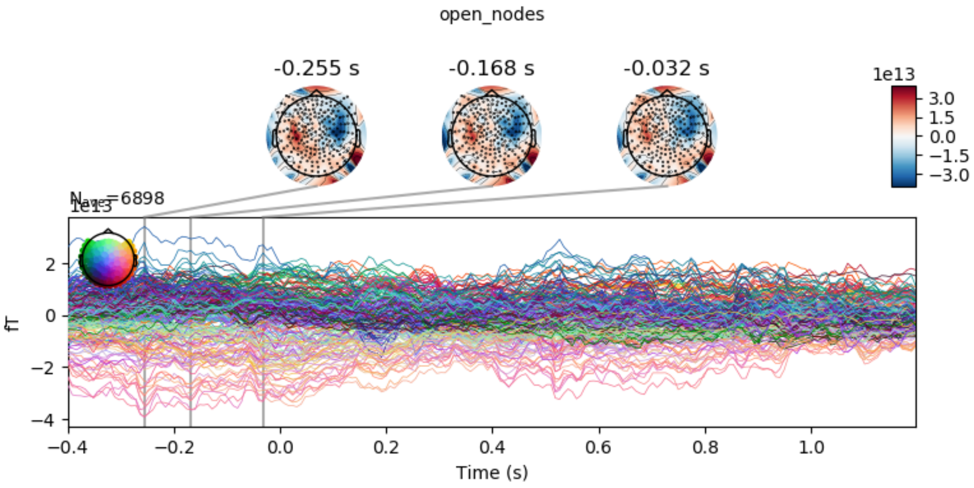


Figure 35: **Open Nodes sensor encoding.** x-axis represents time in seconds relative to word offset; y-axis represents the femto-tesla activity strength of each sensor.

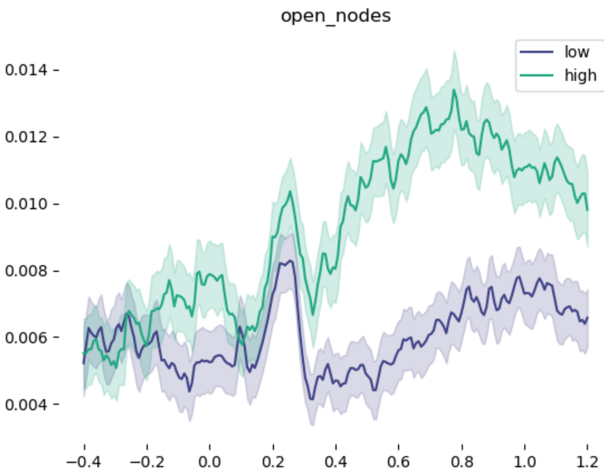


Figure 36: **Open Nodes RMS median split.** x-axis represents time in seconds relative to word offset; y-axis represents the root-mean-square of femto-tesla activity strength over all sensors.

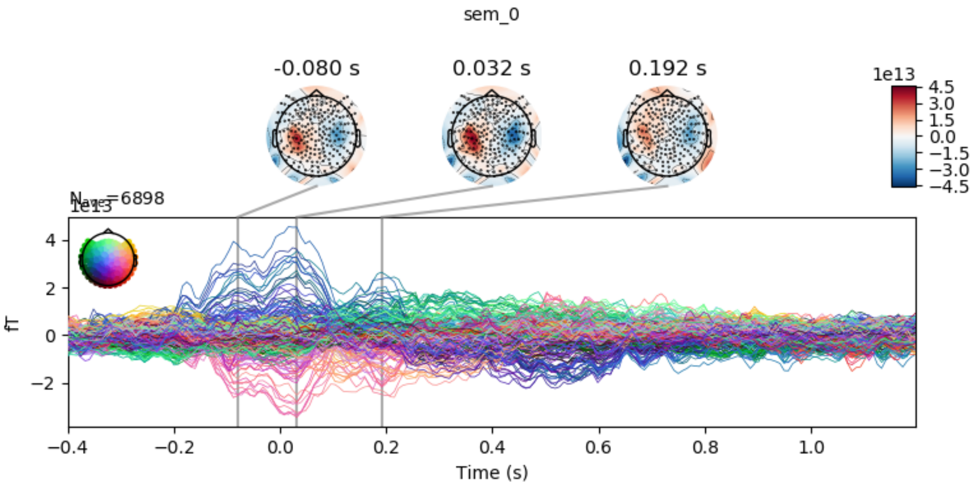


Figure 37: **GloVe PC1 sensor encoding.** x-axis represents time in seconds relative to word offset; y-axis represents the femto-tesla activity strength of each sensor.

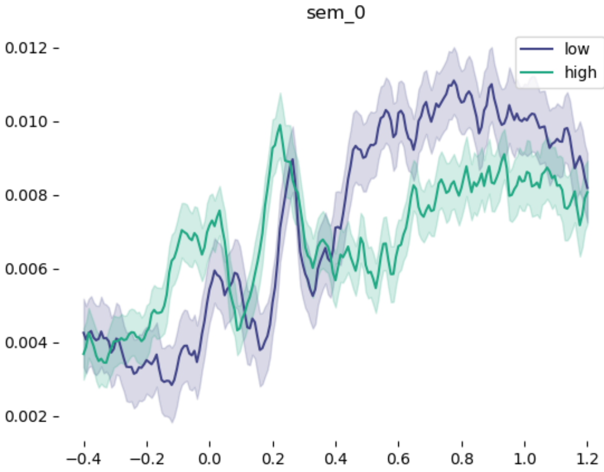


Figure 38: **GloVe PC1 RMS median split.** x-axis represents time in seconds relative to word offset; y-axis represents the root-mean-square of femto-tesla activity strength over all sensors.

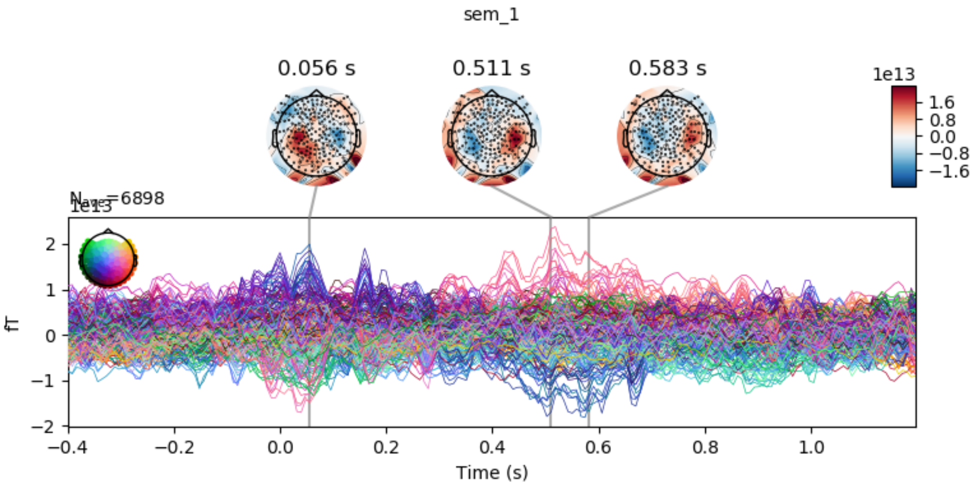


Figure 39: **GloVe PC2 sensor encoding.** x-axis represents time in seconds relative to word offset; y-axis represents the femto-tesla activity strength of each sensor.

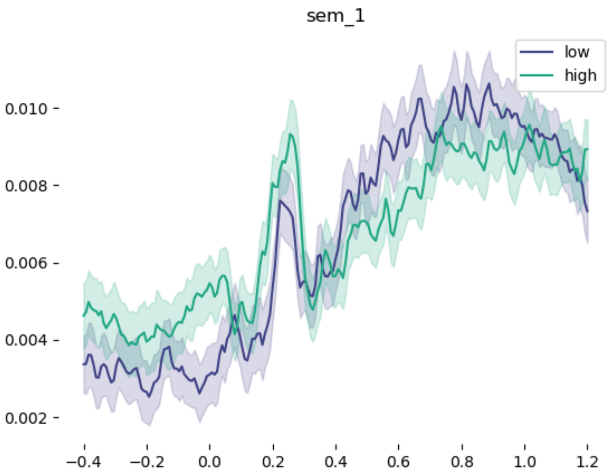


Figure 40: **GloVe PC2 RMS median split.** x-axis represents time in seconds relative to word offset; y-axis represents the root-mean-square of femto-tesla activity strength over all sensors.

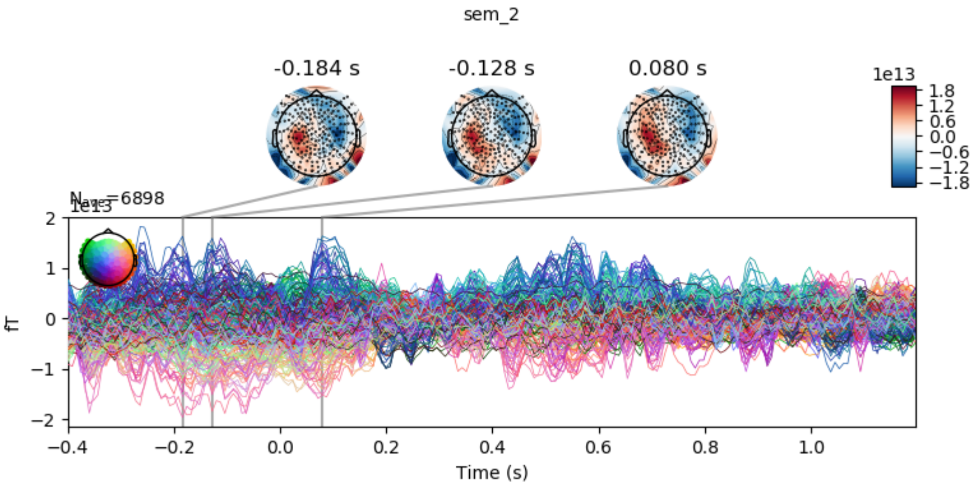


Figure 41: **GloVe PC3 sensor encoding.** x-axis represents time in seconds relative to word offset; y-axis represents the femto-tesla activity strength of each sensor.

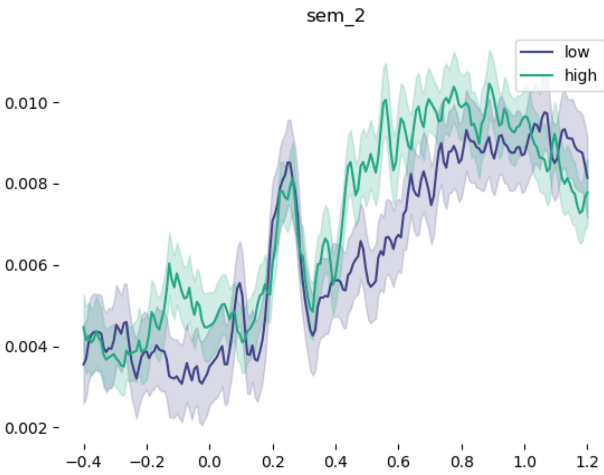


Figure 42: **GloVe PC3 RMS median split.** x-axis represents time in seconds relative to word offset; y-axis represents the root-mean-square of femto-tesla activity strength over all sensors.

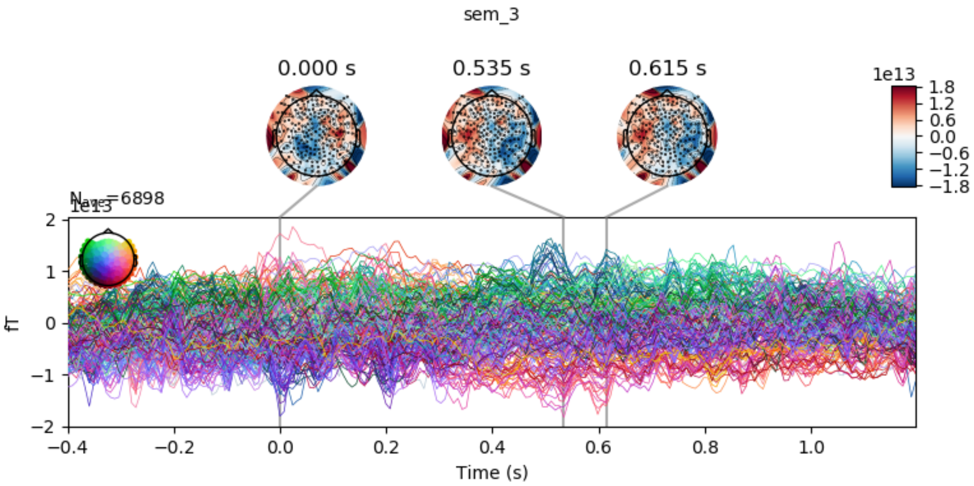


Figure 43: **GloVe PC4 sensor encoding.** x-axis represents time in seconds relative to word offset; y-axis represents the femto-tesla activity strength of each sensor.

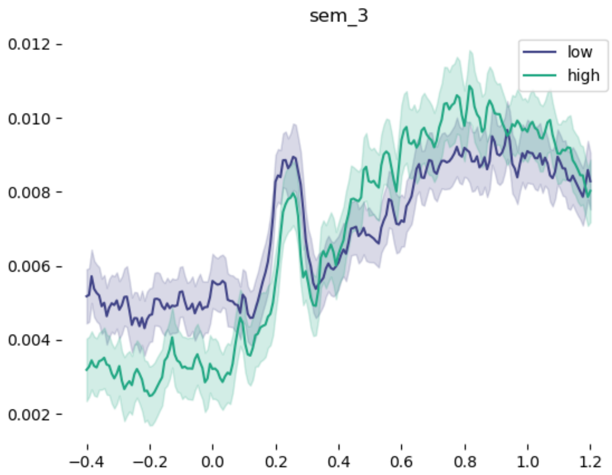


Figure 44: **GloVe PC4 RMS median split.** x-axis represents time in seconds relative to word offset; y-axis represents the root-mean-square of femto-tesla activity strength over all sensors.

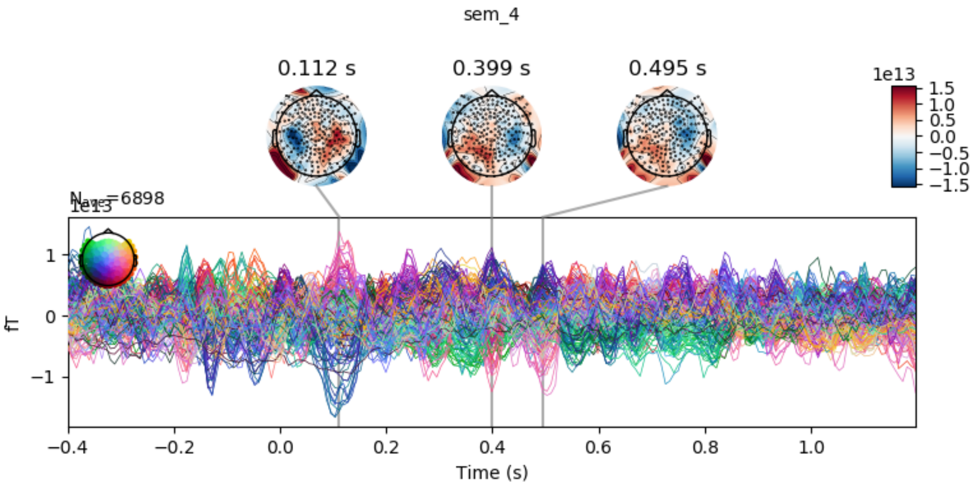


Figure 45: **GloVe PC5 sensor encoding.** x-axis represents time in seconds relative to word offset; y-axis represents the femto-tesla activity strength of each sensor.

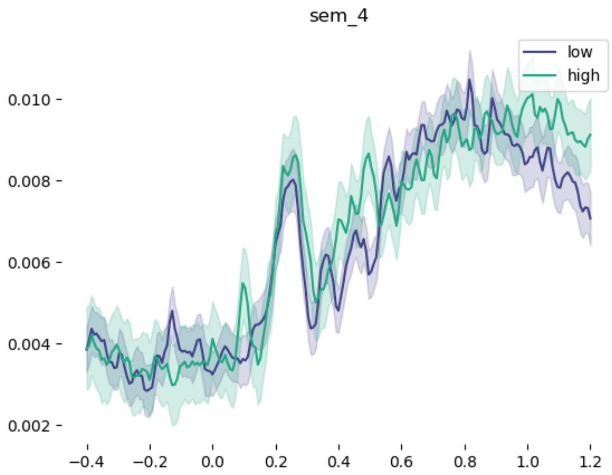


Figure 46: **GloVe PC5 RMS median split.** x-axis represents time in seconds relative to word offset; y-axis represents the root-mean-square of femto-tesla activity strength over all sensors.

| Feature | $\hat{t}$ | $p$ | Window |
|---|---|---|---|
| Approximant | all p > .1 | | |
| Fricative | 2.78 | 0.0245 | 39:168 |
| Nasal | 3.91 | 0.0065 | -16:152 |
| Plosive | all p > .1 | | |
| Vowel | -2.65 | 0.0917 | 1032:1088 |
| Voicing | -2.34 | 0.0981 | -392:-328 |
| Coronal | None | | |
| Dental | all p > .1 | | |
| High Vowel | all p > .1 | | |
| Labial | all p > .1 | | |
| Middle Vowel | None | | |
| Velar | all p > .1 | | |
| Low Vowel | all p > .1 | | |

**Supplementary Table 1.** Results of temporal permutation cluster test applied to the phonetic features relative to word offset.

| Feature | $\hat{t}$ | $p$ | Window |
|---|---|---|---|
| Word Frequency | 4.25 | 0.0001 | -104:632 |
| Word Frequency | -3.18 | 0.0136 | -392:-224 |
| No. Morphemes | 3.56 | 0.0045 | -72:111 |
| No. Morphemes | 3.1 | 0.0649 | 303:360 |
| No. Phonemes | 4.18 | 0.0001 | -136:640 |
| No. PhonSyll | 2.63 | 0.0815 | -128:-64 |
| No. PhonSyll | 2.67 | 0.016 | 303:496 |
| No. Syllables | 4.03 | 0.0061 | -88:120 |
| No. Syllables | 2.9 | 0.0271 | 223:368 |
| Phon. Neigh. Density | 3.2 | 0.0008 | -136:496 |
| Phon. Neigh. Density | 2.37 | 0.0432 | 568:680 |
| Phon. Neigh. Density | 2.79 | 0.0079 | 704:968 |

**Supplementary Table 2.** Results of temporal permutation cluster test applied to the sub-lexical features relative to word offset.

| Feature | $\hat{t}$ | $p$ | Window |
|---|---|---|---|
| Adjective | all p > .1 | | |
| Coord.Conj | 3.34 | 0.0125 | 31:176 |
| Determiner | -2.67 | 0.062 | -288:-224 |
| Determiner | -2.49 | 0.0465 | 680:759 |
| Determiner | -2.9 | 0.0354 | 1120:1200 |
| Noun | 4.88 | 0.0001 | -144:680 |
| Pronoun | -2.71 | 0.0509 | 544:616 |
| Pronoun | -4.08 | 0.0001 | 640:1200 |
| Preposition | -4.15 | 0.0001 | -400:-16 |
| Preposition | -2.59 | 0.0875 | 31:87 |
| Preposition | -3.54 | 0.025 | 288:368 |
| Preposition | -3.64 | 0.0039 | 1024:1200 |
| Adverb | 2.44 | 0.0638 | -400:-320 |
| Adverb | 3.17 | 0.0015 | -256:183 |
| TO | all p > .1 | | |
| Verb | 2.91 | 0.0055 | -216:23 |
| Verb | -2.85 | 0.0812 | 1000:1056 |
| WH-word | None | | |
| Ex-There | None | | |

**Supplementary Table 3.** Results of temporal permutation cluster test applied to the word class features relative to word offset.

| Feature | $\hat{t}$ | $p$ | Window |
|---|---|---|---|
| No. Closing Nodes | 5.82 | 0.0001 | -168:1200 |
| No. Opening Nodes | 3.47 | 0.0001 | -176:528 |
| Sentence End | 6.22 | 0.0001 | -256:1200 |

**Supplementary Table 4.** Results of temporal permutation cluster test applied to the syntactic operation features relative to word offset.

| Feature | $\hat{t}$ | $p$ | Window |
|---|---|---|---|
| Tree Depth | 11.16 | 0.0001 | -400:1200 |
| Tree Depth -1 | 9.63 | 0.0001 | -400:1200 |
| Tree Depth +1 | 10.91 | 0.0001 | -400:1200 |
| No. Open Nodes | 10.52 | 0.0001 | -400:1200 |
| Linear Order | 11.74 | 0.0001 | -400:1200 |
| Distance from end | 10.19 | 0.0001 | -400:1200 |
| Linear Order -1 | 9.64 | 0.0001 | -400:1200 |
| Linear Order +1 | 9.85 | 0.0001 | -400:1200 |
| Distance from start | 10.93 | 0.0001 | -400:1200 |

**Supplementary Table 5.** Results of temporal permutation cluster test applied to the syntactic state features relative to word offset.

| Feature | $\hat{t}$ | $p$ | Window |
|---|---|---|---|
| GloVe PC1 | 5.91 | 0.0001 | -400:1200 |
| GloVe PC2 | 3.45 | 0.001 | -176:632 |
| GloVe PC3 | 2.82 | 0.0625 | -320:-248 |
| GloVe PC3 | 2.86 | 0.0095 | -200:136 |
| GloVe PC3 | 2.63 | 0.0145 | 352:632 |
| GloVe PC3 | 2.31 | 0.085 | 672:736 |
| GloVe PC3 | 2.79 | 0.0064 | 775:1200 |
| GloVe PC4 | 2.54 | 0.0895 | -368:-304 |
| GloVe PC4 | 2.96 | 0.0343 | -264:-111 |
| GloVe PC4 | 3.02 | 0.0194 | 111:352 |
| GloVe PC4 | 2.6 | 0.0666 | 368:455 |
| GloVe PC4 | 2.62 | 0.0412 | 472:616 |
| GloVe PC4 | 3.23 | 0.0054 | 632:1200 |
| GloVe PC5 | 2.7 | 0.0859 | 55:111 |
| GloVe PC5 | 2.7 | 0.0281 | 256:376 |
| GloVe PC5 | 2.57 | 0.0661 | 463:536 |
| GloVe PC6 | 2.89 | 0.0142 | -400:-192 |
| GloVe PC6 | 2.63 | 0.0931 | 0:55 |
| GloVe PC6 | 3.27 | 0.0008 | 128:904 |
| GloVe PC7 | 2.97 | 0.0881 | -400:-344 |
| GloVe PC7 | 3.18 | 0.0004 | 215:1192 |
| GloVe PC8 | 3.98 | 0.0018 | -144:327 |
| GloVe PC8 | 2.69 | 0.0107 | 384:664 |
| GloVe PC8 | 2.66 | 0.0269 | 872:1040 |
| GloVe PC9 | all p > .1 | | |
| GloVe PC10 | all p > .1 | | |

**Supplementary Table 6.** Results of temporal permutation cluster test applied to the semantic word embedding features relative to word offset.

#### *1.8. Replication of reverse hierarchy across sessions*

In order to assess the reliability of our results, we re-ran the hierarchical analysis on just the first and on just the second session of data separately (Figure 4). We find that the results are very similar across sessions, yielding a significant correlation between the averaging decoding timecourse of sessions 1 and 2 (Phonetic: $r = 0.68$, $p < .001$; Sub-lexical: $r = 0.86$, $p < .001$; Lexical: $r = 0.89$, $p < .001$; Syntactic operation: $r = 0.86$, $p < .001$; Syntactic state: $r = 0.62$, $p < .001$; Semantic: $r = 0.68$, $p < .001$).

| Feature | $\hat{t}$ | $p$ | Window |
|---|---|---|---|
| Approximant | | all p > .1 | |
| Fricative | -2.52 | 0.0473 | 1072:1160 |
| Nasal | -3.09 | 0.0522 | 600:672 |
| Nasal | -2.49 | 0.095 | 688:752 |
| Nasal | -2.77 | 0.0118 | 767:928 |
| Plosive | -2.54 | 0.0358 | -400:-304 |
| Plosive | -2.85 | 0.0236 | -288:-184 |
| Plosive | -3.13 | 0.0649 | -168:-111 |
| Plosive | -3.29 | 0.015 | -96:16 |
| Plosive | -2.54 | 0.0225 | 832:952 |
| Vowel | 3.08 | 0.0363 | 111:199 |
| Vowel | 2.88 | 0.0658 | 215:280 |
| Voicing | 3.33 | 0.0378 | 183:280 |
| Voicing | 2.7 | 0.0379 | 327:447 |
| Coronal | 3.36 | 0.0881 | 248:296 |
| Dental | 3.23 | 0.0873 | 136:183 |
| Glottal | 2.62 | 0.0635 | -88:0 |
| Glottal | 2.83 | 0.0106 | 64:424 |
| Glottal | 2.34 | 0.0932 | 504:568 |
| Glottal | 2.58 | 0.0954 | 584:640 |
| Glottal | 2.61 | 0.0776 | 712:783 |
| Glottal | 2.31 | 0.0952 | 936:1000 |
| High Vowel | 2.97 | 0.0492 | 136:199 |
| Labial | | all p > .1 | |
| Middle Vowel | -2.51 | 0.0398 | -368:-288 |
| Velar | | all p > .1 | |
| Low Vowel | 3.06 | 0.0595 | 231:288 |

**Supplementary Table 7.** Results of temporal permutation cluster test applied to the phonetic features relative to word onset.

| Feature | $\hat{t}$ | $p$ | Window |
|---|---|---|---|
| Word Frequency | 3.44 | 0.0107 | 191:303 |
| Word Frequency | -2.79 | 0.0947 | -176:-128 |
| No. Morphemes | | all p > .1 | |
| No. Phonemes | | all p > .1 | |
| No. PhonSyll | 2.9 | 0.0717 | -360:-296 |
| No. PhonSyll | 2.99 | 0.0627 | -24:47 |
| No. PhonSyll | 3.08 | 0.0412 | 136:240 |
| No. PhonSyll | 2.58 | 0.029 | 256:424 |
| No. PhonSyll | 2.7 | 0.0018 | 472:1080 |
| No. PhonSyll | 2.66 | 0.0546 | 1104:1200 |
| No. Syllables | | None | |
| Phon. Neigh. Density | 2.59 | 0.0263 | 111:248 |
| Phon. Neigh. Density | 2.36 | 0.0594 | 368:463 |

**Supplementary Table 8.** Results of temporal permutation cluster test applied to the sub lexical features relative to word onset.

| Feature | $\hat{t}$ | $p$ | Window |
|---|---|---|---|
| Adjective | -2.65 | 0.0214 | -160:-16 |
| Coord.Conj | 2.49 | 0.0896 | -16:47 |
| Coord.Conj | 2.98 | 0.0381 | 64:152 |
| Coord.Conj | 3.35 | 0.0075 | 199:360 |
| Determiner | | all p > .1 | |
| Noun | 3.32 | 0.0291 | 215:327 |
| Noun | 3.41 | 0.007 | 416:632 |
| Noun | 2.59 | 0.0446 | 648:759 |
| Pronoun | -3.32 | 0.0753 | 656:704 |
| Pronoun | -3.63 | 0.0008 | 791:1200 |
| Preposition | -3.43 | 0.0005 | -400:-32 |
| Adverb | 2.38 | 0.088 | -352:-288 |
| Adverb | 2.5 | 0.0598 | 136:215 |
| Adverb | 3.03 | 0.0525 | 303:376 |
| Adverb | 3.25 | 0.0007 | 416:767 |
| Adverb | 2.49 | 0.0961 | 848:904 |
| Adverb | 3.07 | 0.0709 | 1048:1104 |
| TO | | all p > .1 | |
| Verb | 2.97 | 0.0672 | 64:128 |
| Verb | 3.19 | 0.045 | 344:424 |
| WH-word | | all p > .1 | |
| Ex-There | | None | |

**Supplementary Table 9.** Results of temporal permutation cluster test applied to the word class features relative to word onset.

| Feature | $\hat{t}$ | $p$ | Window |
|---|---|---|---|
| No. Closing Nodes | 3.53 | 0.0144 | 248:392 |
| No. Closing Nodes | 4.09 | 0.0004 | 408:696 |
| No. Closing Nodes | 3.7 | 0.0001 | 712:1176 |
| No. Closing Nodes | -4.43 | 0.0002 | -400:-72 |
| No. Closing Nodes | -3.4 | 0.026 | -55:47 |
| No. Opening Nodes | 2.41 | 0.0787 | 95:168 |
| No. Opening Nodes | 2.77 | 0.0543 | 256:335 |
| No. Opening Nodes | 3.14 | 0.0109 | 376:536 |
| No. Opening Nodes | 2.81 | 0.0876 | 552:608 |
| Sentence End | 4.08 | 0.0001 | 248:1128 |
| Sentence End | 4.4 | 0.0579 | 1144:1200 |
| Sentence End | -2.79 | 0.0852 | -312:-248 |

**Supplementary Table 10.** Results of temporal permutation cluster test applied to the syntactic operation features relative to word onset.

| Feature | $\hat{t}$ | $p$ | Window |
|---|---|---|---|
| Tree Depth | 11.17 | 0.0002 | -400:47 |
| Tree Depth | 10.06 | 0.0001 | 72:1200 |
| Tree Depth -1 | 10.12 | 0.0005 | -400:-40 |
| Tree Depth -1 | 9.28 | 0.045 | -24:47 |
| Tree Depth -1 | 8.6 | 0.0001 | 64:1152 |
| Tree Depth -1 | 8.15 | 0.0587 | 1168:1200 |
| Tree Depth +1 | 10.72 | 0.0001 | -400:47 |
| Tree Depth +1 | 9.39 | 0.0001 | 64:1200 |
| No. Open Nodes | 10.65 | 0.0002 | -400:47 |
| No. Open Nodes | 9.24 | 0.0001 | 64:1200 |
| Linear Order | 13.24 | 0.0001 | -400:47 |
| Linear Order | 11.29 | 0.0001 | 64:1200 |
| Distance from end | 11.82 | 0.0313 | -400:-280 |
| Distance from end | 10.91 | 0.0036 | -264:0 |
| Distance from end | 10.9 | 0.055 | 16:47 |
| Distance from end | 9.96 | 0.0174 | 64:288 |
| Distance from end | 9.65 | 0.0001 | 303:1200 |
| Linear Order -1 | 10.69 | 0.0002 | -400:47 |
| Linear Order -1 | 9.15 | 0.0001 | 64:1200 |
| Linear Order +1 | 11.02 | 0.0002 | -400:47 |
| Linear Order +1 | 9.92 | 0.0001 | 64:1200 |
| Distance from start | 12.07 | 0.0002 | -400:47 |
| Distance from start | 10.92 | 0.0001 | 64:1200 |

**Supplementary Table 11.** Results of temporal permutation cluster test applied to the syntactic state features relative to word onset.

| Feature | $\hat{t}$ | $p$ | Window |
|---|---|---|---|
| GloVe PC1 | 2.93 | 0.0063 | -400:16 |
| GloVe PC1 | 4.66 | 0.0019 | 64:472 |
| GloVe PC1 | 4.47 | 0.0001 | 488:1200 |
| GloVe PC2 | 3.2 | 0.0231 | -400:-240 |
| GloVe PC2 | 3.43 | 0.0087 | -224:47 |
| GloVe PC2 | 3.13 | 0.0921 | 64:111 |
| GloVe PC2 | 3.65 | 0.0002 | 136:968 |
| GloVe PC3 | 2.62 | 0.0735 | 280:352 |
| GloVe PC3 | 2.73 | 0.0061 | 744:1200 |
| GloVe PC4 | 3.15 | 0.0354 | -400:-272 |
| GloVe PC4 | 3.39 | 0.0125 | -200:47 |
| GloVe PC4 | 3.28 | 0.0005 | 64:1200 |
| GloVe PC5 | | all p > .1 | |
| GloVe PC6 | 2.79 | 0.0127 | -400:-128 |
| GloVe PC6 | 2.4 | 0.0779 | 79:152 |
| GloVe PC6 | 2.36 | 0.0421 | 696:824 |
| GloVe PC6 | 2.6 | 0.04 | 1000:1120 |
| GloVe PC7 | 2.54 | 0.0821 | 791:872 |
| GloVe PC7 | 2.99 | 0.0115 | 888:1200 |
| GloVe PC8 | 3.6 | 0.0021 | 103:376 |
| GloVe PC9 | | all p > .1 | |
| GloVe PC10 | | all p > .1 | |

**Supplementary Table 12.** Results of temporal permutation cluster test applied to the semantic word embedding features relative to word onset.
